# Climbing fibers encode the gradient of a loss function for the cerebellum

**DOI:** 10.64898/2026.07.27.741034

**Authors:** Jafar Doostmohammadi, Nazanin Mohammadrezaei, Hisham Y. Elseweifi, Alana D. Chandler, Elijah A. Taeckens, Reza Shadmehr

## Abstract

Neurons in the brain are often many synapses away from motoneurons, yet if a movement results in error, each distant neuron needs a teacher that considers its specific contribution to production of that movement. This credit assignment problem is solved in machine learning via gradient descent of a loss function, where the loss defines the subjective cost incurred by error. Does the brain use gradient descent to teach individual neurons? We trained marmosets to make saccades to visual targets and then varied the loss by assigning reward value to each target. The climbing fibers, which are the teachers of Purkinje cells (P-cells) in the cerebellum, used a multiplicative encoding to scale the spatial properties of the error vector with its reward properties, incorporating reward prediction errors. Using spike-triggered suppression of P-cells, we quantified the potent vector that mapped each P-cell’s output to eye movements and discovered that the climbing fibers were not merely transmitting errors. Rather, they were providing a signal that was, on average, proportional to the dot product of the reward dependent error vector upon the P-cell’s potent vector. Thus, the climbing fibers solved the credit assignment problem by providing the gradient of a loss function with respect to the output of their individual P-cells.

---

Learning aims to minimize loss, where loss is a scalar function that provides a metric for how well an action is performed ^1^. For example, during tennis if a serve lands outside the lines, the loss associated with that error is larger at match point than during practice (Fig. 1A). This implies that when we perform a movement and experience error, the loss, and the resulting learning, should depend on both the spatial properties of the error vector and the reward context in which that error was experienced. Indeed, learning from error can be modulated by reward ^2–8^. This hints that the brain has access to a teaching signal that combines sensory properties of error with reward.

**Fig. 1.**
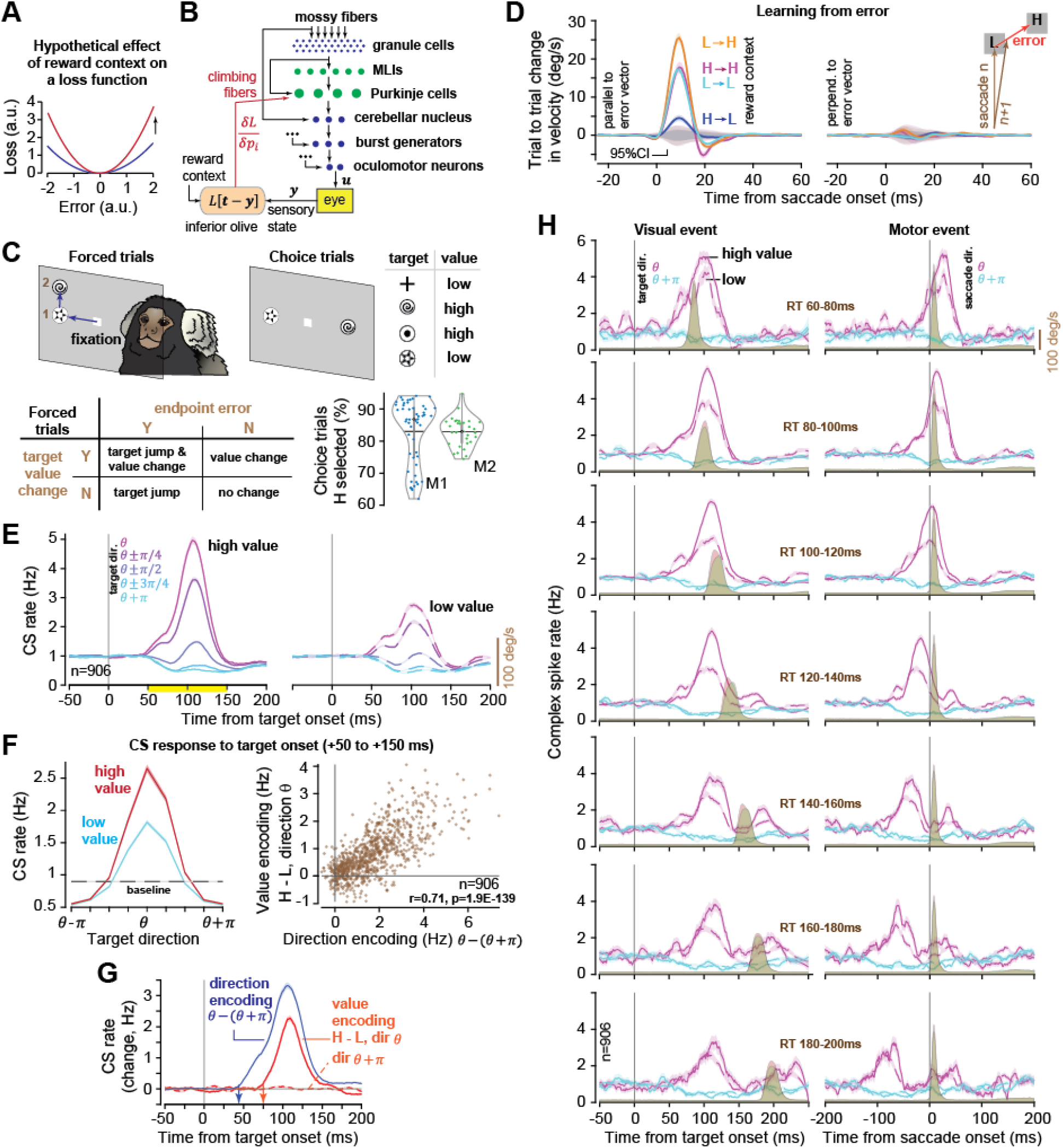
Climbing fibers multiplicatively encoded the direction of a random saccade target and its value. **A**. Hypothetical loss function, illustrating that loss typically grows with error magnitude, but depends on the reward context. **B**. When a saccade ends in error ***e*** = ***t*** − ***y***, the loss associated with that error must be transmitted to the P-cells. The hypothesis is that the climbing fibers solve the credit assignment problem by transmitting the gradient of the loss with respect to the output *p_i_* of each P-cell. **C**. Marmosets were trained to associate visual stimuli with high or low amounts of reward. In forced trials, during the primary saccade sometimes the target jumped, inducing an endpoint error, and/or changed value, inducing a reward prediction error. Probability of selecting the high valued option is shown for choice trials (each dot is a recording session). **D**. Learning from error. Trial-to-trial change in saccade trajectory, as measured via the time series of velocity vectors projected onto unit vectors, one parallel and the other perpendicular to the endpoint error vector. Gray region indicates 95% confidence interval. **E**. CS firing rates as a function of the direction and value of the primary target. Preferred direction is labeled as *θ*. **F**. Left: CS rates as a function of target direction and target value. Right: Strength of value coding vs. direction coding for each climbing fiber. **G**. Timing of reporting target direction (CS rates in direction *θ* vs. *θ* + *π*), and target value (CS rates for high vs. low valued target, direction *θ* or *θ* + *π*). **H**. CS rates for high and low valued targets binned by reaction time. Error bars are SEM.

In machine learning, backpropagation provides a teaching signal that conveys the gradient of the loss for each neuron ^9^. This gradient solves the credit assignment problem by combining steepness of the loss with respect to error, which depends on the reward context, with the strength of each neuron’s contribution to production of error, which depends on the neuron’s downstream influence on the movement. The success of backpropagation suggests that a good teacher cannot simply broadcast errors ^10^. Rather, it must first combine the sensory properties of error with its reward context ^2^, then tailor this information to match each neuron’s unique contribution to production of error ^11^.

The cerebellum is an attractive substrate for understanding how the brain learns to minimize loss functions. Its principal neurons, Purkinje cells (P-cells), are many synapses away from motoneurons (Fig. 1B), yet their plasticity is required for most forms of motor learning ^12,13^. Indeed, error driven changes in P-cell output can be a predictor of trial-by-trial changes in behavior ^14–16^. Notably, P-cell plasticity depends on a teacher whose activities can be measured via its climbing fiber input ^16,17^. Because climbing fibers also convey reward information to the cerebellum ^18–21^, we wondered whether their activities could be understood in terms of the gradient of a loss function.

We trained marmosets to associate visual targets with various amounts of reward and then as they made saccades, we manipulated both the endpoint errors and the reward context. The climbing fiber input to each P-cell peaked for a specific direction of error, with a firing rate that increased with the magnitude of error. Notably, this relationship was multiplicatively modulated by reward. Next, we used spike-triggered suppression of P-cells to measure the resulting eye displacement, i.e., the potent vector ^22,23^, and found that the climbing fiber input was, on average, proportional to the dot product of the reward dependent error vector and the P-cell’s potent vector. Thus, climbing fibers provided the gradient of a loss function.

## The requirements of a good teacher

To conceptualize the credit assignment problem and how gradient descent solves it, consider a saccade that moves the eyes toward a target location ***t***. The motoneurons that innervate the extraocular muscles produce their output due to the inputs that they receive from the brainstem burst generators, which in turn receive inputs from the superior colliculus, omnipause neurons, and the fastigial nucleus of the cerebellum (Fig. 1B). The fastigial nucleus receives its inputs from mossy fibers and P-cells. When the saccade ends, the difference between the target ***t*** and eye position ***y*** is an error vector, ***e*** = ***t*** − ***y***. Let us assume that the brain evaluates this error ^1^ by a loss function *L*(***e***), producing a scalar quantity that, analogous to a tennis serve at match point, depends on the subjective context of the error ^4,24^.

The cerebellum’s integrity is required for reducing saccade errors ^25–27^, and the climbing fibers in the oculomotor region of the vermis encode these errors ^28–30^, generating complex spikes (CS) that induce long-term depression at coactive parallel fiber – Purkinje cell synapses ^31^. This CS-induced plasticity produces trial-to-trial change in the P-cell’s output ^13^ (i.e., simple spikes), which in turn produces trial-to-trial change in eye trajectory ^13–15^. Gradient descent requires that in response to error, the synapse of granule cell *j* onto Purkinje cell *i* be updated in proportion to the gradient of the loss function with respect to the synaptic strength *w_ji_*:

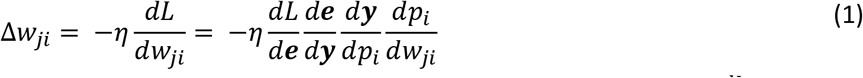

This equation contains two vector quantities: the gradient of the loss with respect to error 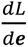, and the gradient of eye position with respect to simple spikes 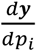 (note that 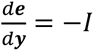, the identity matrix.). The term 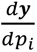 is the potent vector of P-cell *i* because it describes the eye displacement that takes place when this P-cell modulates its firing rate ^22^. The term 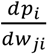 represents the change in P-cell’s simple spike output *p_i_* due to a change in synaptic weight *w_ji_*. If we approximate the activity of the P-cell as a linear sum of its parallel fiber inputs, *p_i_* (*t*) ∝ ∑_*j*_*w_ji_g_j_*(*t*), then 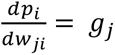, the activity of granule cell *j*. Thus, the weight update equation becomes:

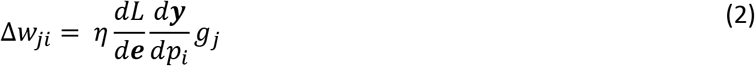

Parallel fiber to P-cell plasticity results from coactivity of the climbing fiber input and the parallel fiber synapse that needs to be modified ^31^. This learning rule can be implemented if the climbing fiber encodes the projection of the loss gradient with respect to error, 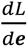, onto the P-cell’s potent vector, 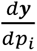 since the multiplication by *g_j_* is already supplied by the requirements of synaptic coactivation. Thus, gradient descent implies that the climbing fiber input to P-cell *i*, written as *c_i_*, should represent the following:

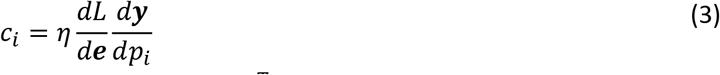

Let us represent the potent vector of P-cell *i* via 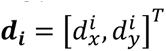, where 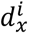 and 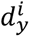 are the displacements in the horizontal and vertical components of eye movements. To consider the effects of reward, suppose that our loss function is a reward dependent quadratic function of error 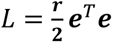, where *r* is a scalar indicating the reward context. In that case, we have:

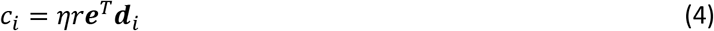

Eq. (4) states that the climbing fiber input to P-cell *i* should exhibit directional tuning with respect to the error vector: the CS rate will be maximum when ***e*** is parallel with the potent vector ***d*** of the P-cell, but zero (or at baseline) when the two vectors are perpendicular. It also specifies that reward should multiplicatively scale this directional tuning. Do climbing fibers provide a signal that obeys the teacher-student requirements of gradient descent?

## Learning was modulated by the reward context

We trained two marmoset monkeys to associate four visual targets with low or high amounts of reward (Fig. S1). To check that they understood this relationship, in choice trials (20% probability) we presented two options, equidistant from center fixation along a random axis (Fig. 1C). The subjects selected the high valued target on 83±0.9% of the trials (mean±SEM).

To induce endpoint error, in forced trials (80% probability) we presented a single target at a random direction but then on some trials (Fig. S1C) jumped its position mid-saccade to another random position. To manipulate the reward context, in a fraction of trials (Fig. S1C) we jumped the target and changed its value. For example, if the primary target was high valued, but the secondary target was low valued, completion of the trial resulted in both a sensory prediction error (i.e. target is not on the fovea), and a reward prediction error (i.e., high reward was promised but low reward was earned). The result was a 2×2 task in which the subjects experienced both types of errors (Fig. 1C).

On each forced trial *n* we measured endpoint error via the vector ***e****_n_*. This vector connected the end of the primary saccade to the center of the secondary target (Fig. 1D). We measured learning by comparing the saccade trajectory ***ẏ***(*t*), i.e., the two-dimensional eye velocity vector as a function of time, in consecutive (*n* and *n* + 1) or nearly consecutive (*n* and *n* + 2) primary saccades in which the primary targets were identical in both position and value. We projected Δ***ẏ***(*t*) upon a pair of unit vectors, one parallel and the other perpendicular to ***e****_n_*. The change in trajectory Δ***ẏ***(*t*) was in the direction of ***e****_n_*, with little or no component perpendicular to it (Fig. 1D). Thus, the subjects learned from error.

However, the amount of learning depended on the reward context. Learning was largest when the primary target was low valued but then changed to high (LH condition, Fig. 1D), smaller if the target value did not change (HH and LL conditions), and smallest when the primary target was high valued but then changed to low (HL condition) (linear mixed effects, M1: F(1,240)=49.0, p=1.6E-11; M2: F(1,240)=38.37, p=2.5E-9). These behavioral results were consistent in both monkeys (Fig. S2).

In summary, target jumps induced endpoint errors, and this promoted trial-to-trial learning. However, the amount of learning depended on the reward context. Learning was largest in the LH context (RPE+, reward earned was more than promised by the primary target), smallest in the HL context (RPE-, reward earned was less than promised).

## Climbing fibers reported both the direction of the visual event and its reward value

We used silicon probes to record from 906 definitive P-cells in lobule VI and VII of the cerebellar vermis. Our dataset included 220 P-cells with both simple and complex spikes (SS and CS), and 686 P-cells with only complex spikes (Fig. S3).

Trials began with fixation of a center target. In forced trials, when a primary target appeared in a random direction, the climbing fibers responded with a large increase in CS rates if the target was in a preferred direction, labeled as *θ*, and a suppression below baseline when it was in direction *θ* + *π*. Across the P-cells, the distribution of *θ* spanned the entire visual field (Fig. S4), though in our sample there was a clustering toward the left side. Notably, the response in direction *θ* was much stronger when the target had high value (Fig. 1E, CS rate 50 to 150ms post visual onset, high vs. low value, paired t-test, t(905) = 28.97, p<0.001). Consequently, the CS rates presented a tuning function with respect to the direction of the target ^30,32^, but target value multiplicatively modulated this tuning, producing the largest increase when the target was in direction *θ*, but little or no change when target was in direction *θ* + *π* (Fig. 1F, left subplot).

These results quantified the average response across the P-cells, leaving open the question of whether some climbing fibers responded primarily to target value, while others responded to target direction. To measure encoding of target value, in each P-cell we quantified the difference in CS rates between high and low-reward targets in direction *θ* (individual cells, Fig. S5). To measure encoding of target direction, we measured the difference in CS rates between directions *θ* and *θ* + *π*. This within-cell analysis revealed a very strong correlation between encoding of target value and target direction (Fig. 1F, right panel, r=0.71, p=1.9E-139). A similar pattern was present when the CS response was aligned to saccade onset (Fig. S5). Thus, climbing fibers that strongly encoded the direction of the target also strongly reported the value of that target.

However, the direction and value information were encoded at slightly different times. To estimate the timing of direction encoding, in each P-cell we compared the CS rates in directions *θ* and *θ* + *π* and found that on average, the rates climbed above chance at 43±0.3ms after target onset (mean±SEM, Fig. 1G, blue trace). To estimate the timing of value encoding, in each P-cell we compared the CS rates between high and low-reward targets in direction *θ* and found that the rates rose above chance at 68.7±1.6ms after target onset (Fig. 1G, solid orange trace). In contrast, the value information was completely missing when the target was in direction *θ* + *π* (Fig. 1G, dashed orange trace). Thus, information regarding target value arrived 25ms after the information regarding its direction.

We next asked whether the modulation of the CS rates was principally due to the visual event (target onset), or the motor event (saccade). We binned the trials based on reaction time (RT) and found that when RTs were short, the CS rates peaked after saccade onset (Fig. 1H, 1^st^ row), but when RTs were long, the CS rates peaked before saccade onset (Fig. 1H, last row). Indeed, regardless of RT, the CS rates peaked at around 100ms after target onset. Thus, CS rates were principally modulated by the appearance of the visual stimulus, and to a lesser extent by the movement toward it.

As expected, saccades toward the high valued targets were faster (Fig. S6A, M1, 52 sessions, signrank z=6.27 p=0; M2 31 sessions, z=4.86, p=0), but when we normalized for peak speed, CS rates were nearly twice as large for high valued as compared to low valued targets (Fig. S6B, 2-way RMANOVA, F(1,892)=86, p=7.32E-31). Thus, the climbing fibers primarily encoded target value, and to a lesser extent vigor of the movement.

In summary, at short latency (∼50ms) the climbing fibers reported the direction of the visual target, and then 25ms later they reported its reward value. At 50-150ms post target onset, the value encoding multiplicatively enhanced the direction encoding.

## Climbing fibers reported the subjective value of the option in their visual field

If a goal is deemed valuable, people and monkeys start their saccades earlier ^33–36^. Indeed, in forced trials the RTs were shorter for high valued targets (Fig. 2A, M1, 52 sessions, signrank z=-6.27, p=0, M2, 31 sessions, signrank z=-4.86, p=0). However, even for a constant target value, RTs varied widely (Fig. 1H), reflecting the changing motivational state of the animal. Earlier work had demonstrated that for a constant valued stimulus, as motivation declined, RTs increased while dopamine response to the stimulus decreased ^37^. We wondered whether the climbing fibers exhibited a similar relationship to RT.

**Fig. 2.**
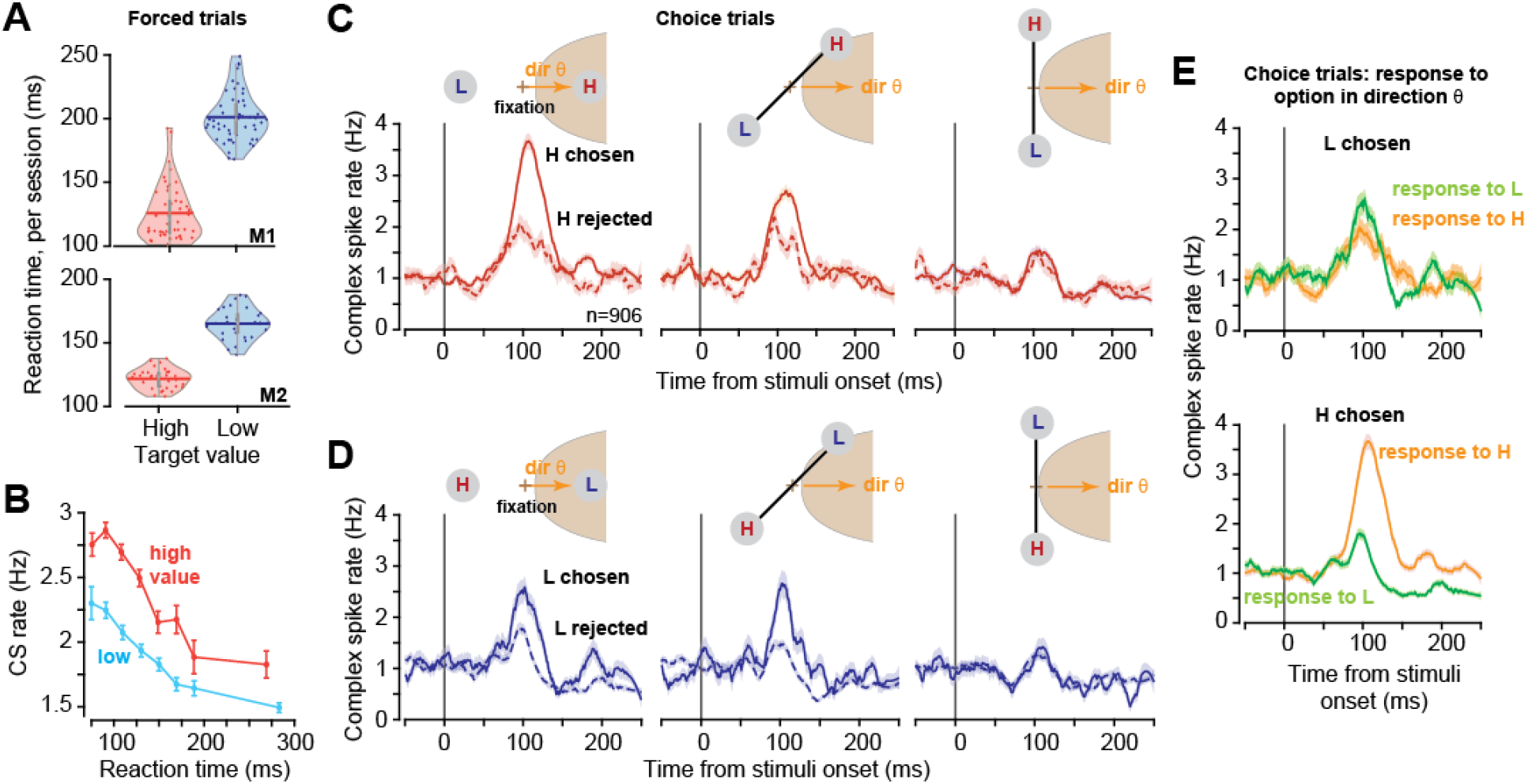
Climbing fibers reported the subjective value of the option in their visual field. **A**. Reaction times in forced trials in each monkey (each dot is a single session). **B**. CS rates as a function of reaction time in response to the primary target in direction *θ* (50-150ms period). As motivation declined, so did the CS response. **C**. CS rates in choice trials conditioned by the location of the high valued stimulus with respect to *θ* and whether that option was selected or rejected. When the high valued stimulus was selected, the CS response to that stimulus was greater than when the same option was rejected. **D**. Similar to part (C), but for low valued stimuli. When the low valued stimulus was selected, the CS response to that stimulus was greater than when the same option was rejected. **E**. Top: CS rates in choice trials in which the low-valued stimulus was selected. Bottom: CS rates in trials in which the high-valued stimulus was selected. Error bars are SEM.

Indeed, for both the high and low valued targets, as RT increased the CS response to the target decreased (Fig. 2A, high valued, Spearman r=-0.95, p=0.0011, low valued, r=-1.0, p=5.0E-5). This raised the possibility that the climbing fibers provided subjective value information to the cerebellum. To explore this idea, we examined choice trials and asked whether during deliberation, the information transmitted by the climbing fibers was, on average, a predictor of the eventual choice.

During choice trials, sometimes the high-reward option appeared in direction *θ* of a P-cell and the subject chose that option, while in other trials the same option appeared in the same location, but it was rejected. The CS response to the high-reward target was greater when it was chosen, as compared to when it was rejected (Fig. 2C, left panel, t(859)=9.69, p=0). Similarly, in trials where the low-reward target appeared in direction *θ* of a P-cell and the subject chose it, the CS response to that target was greater as compared to when it was rejected (Fig. 2D, left panel, t(870)=6.43, p=2E-10). A similar but weaker pattern was present when the targets appeared in direction *θ* ± *π*/4 (middle column of Figs. 2C & 2C), and the pattern disappeared when the targets appeared in direction *θ* ± *π*/2 (right column of Figs. 2C & 2D).

The interesting trials were those in which the subject selected the low reward option. To understand why this was the case, we considered choice trials in which the low reward option appeared in direction *θ* of a P-cell, conditioned on whether that option was selected. Remarkably, when the low-reward option was selected, the CS response to that stimulus was marginally greater as compared to when the high-reward stimulus appeared in the same location but was rejected (Fig. 2E, top panel, t(824)=2.04, p=0.042, 50-120ms period). Similarly, if the high-reward option was selected, the response to that stimulus was greater as compared to when the low-reward stimulus appeared at the same location but was rejected (Fig. 2E, lower panel, t(905)=20.16, p=0).

In summary, as the motivational state of the subject declined, so did the climbing fiber response to the targets. In choice trials, regardless of whether the high or low valued target was chosen, the climbing fiber response to the stimulus in its visual field was greater if that option was eventually chosen. Thus, the climbing fibers appeared to report the subjective value of the stimulus.

## Climbing fibers reported reward prediction errors

Did the climbing fibers convey merely the reward value of the target or were they also sensitive to reward prediction errors (RPEs)? To answer this question, in a fraction of the forced trials (Fig. S1) we changed the value of the target during the primary saccade but kept its position unchanged (Fig. 3A). For example, in LH trials the subject was presented with a low valued target but during the primary saccade the target changed to high value. Thus, the primary saccade was made in anticipation of low reward but after its conclusion, the value changed and the subject subsequently received a high reward. Notably, there were no target jumps in these trials, i.e., RPE without SPE.

**Fig. 3.**
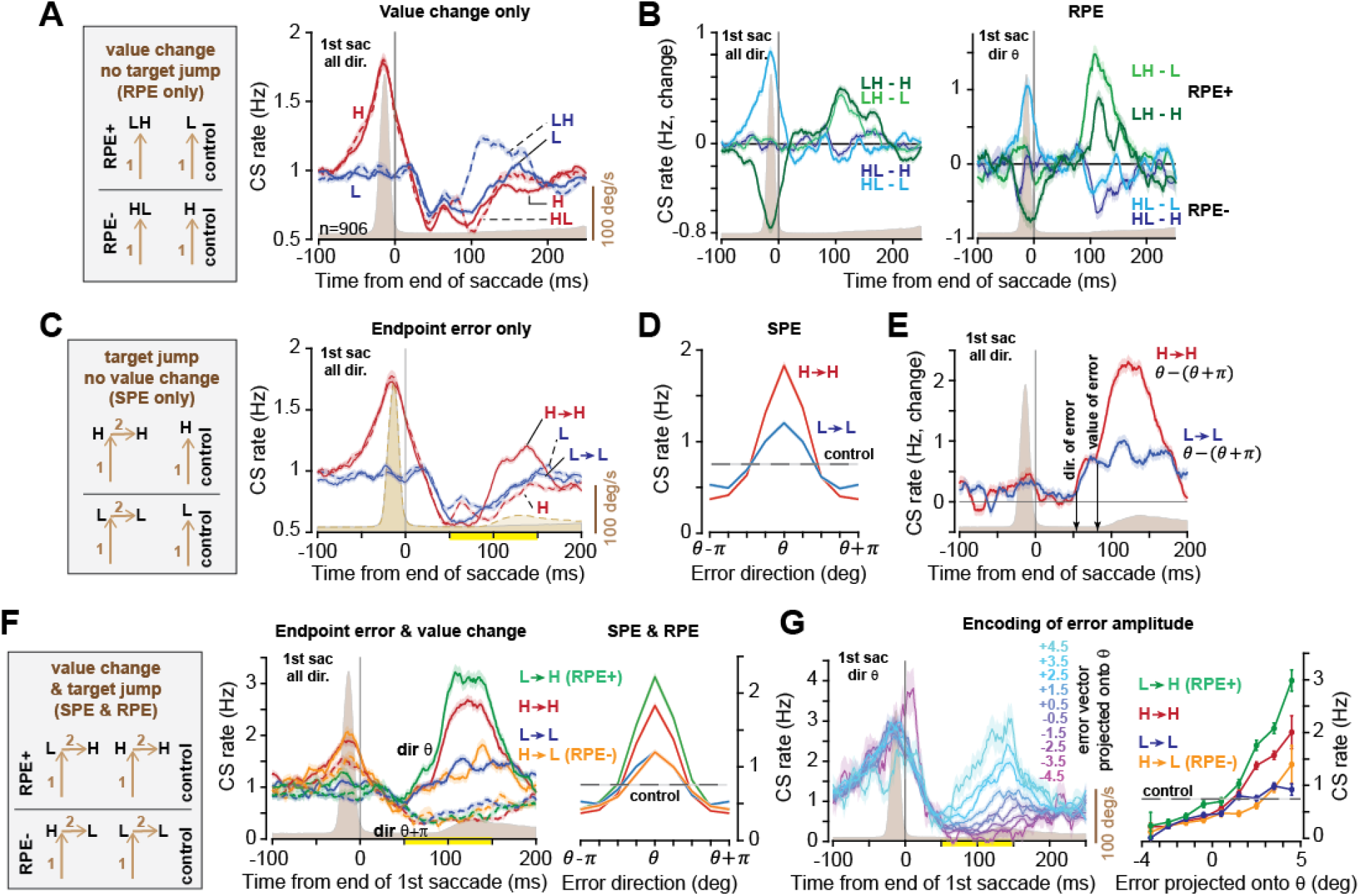
RPE multiplicatively enhanced the climbing fiber response to endpoint error. **A**. RPE only trials: value change without target jump. CS rates are aligned to conclusion of the primary saccade. **B**. Response to RPE+ calculated as the firing rates in LH trials minus control. Response to RPE-calculated as the rate in HL trials minus control. Left plot shows primary saccade in all directions, right plot shows primary saccade in direction *θ*. **C**. SPE only trials: target jump without value change. Response to endpoint error across various reward conditions. **D**. CS response to endpoint error (50-150ms post primary saccade period), for high and low reward conditions. The dashed line shows the response in the control condition (no target jump). **E**. Timing of encoding the direction of error, and the value of error. The traces display change in CS rates when the endpoint error was in direction *θ* vs. *θ* + *π*, for the high and low reward condition. **F**. RPE & SPE trials. Left: CS response to endpoint error in directions *θ* and *θ* + *π* for various reward contexts. Right: CS response to endpoint error (50-150ms post primary saccade period) as a function of direction of error and reward context. Control is same as part (D). **G**. Left: CS rates as a function of the size of the error vector projected upon a unit vector in direction *θ*. Right: CS rates as a function of error size for the various reward contexts. Control is same as part (D). Error bars are SEM.

We compared the LH trials (i.e., value reduced) with L trials (i.e., value not changed) (LH - L) and found that the CS rates rose at around 85ms following conclusion of the primary saccade, peaking at around 110ms (Fig. 3B, left plot). Similarly, when the target was of high value but changed to low value, the CS response in HL vs. H trials declined at around 85ms, exhibiting a minimum at around 110ms (Fig. 2B, left plot, HL - H). These results magnified when the primary saccade was in direction *θ* (Fig. 3B, right plot, HL-H). Thus, when there was a difference between the reward promised and the reward that would be earned, the climbing fibers reported that difference to the cerebellum. This suggested that the climbing fibers may be reporting RPEs.

However, these CS responses may have been merely an encoding of the high (or low) valued stimulus that was in the visual field when the primary saccade ended, not an RPE. To check this, we compared LH trials with H trials. In both LH and H trials, when the primary saccade ended, the stimulus on the screen had high value. However, in one case (LH) the value had increased, whereas in another case (H) it had not changed. We found that the CS rates were greater in LH trials with respect to H trials (Fig. 3B, left plot, LH - H), particularly if the primary saccade was in direction *θ* (Fig. 3B, right plot, LH-H). Similarly, in HL trials as compared to L trials, the CS rates decreased (Fig. 3B, left plot, HL - L), particularly if the primary saccade was in direction *θ* (Fig. 3B, right plot, HL - L). Thus, in addition to reporting the value of the target (Fig. 1F), the CS rates also reported RPEs.

Although there were no target jumps in these trials, there were occasional small, catch-up saccades that occurred at the end of the primary saccades (Fig. S7A). These catch-up saccades may have affected the CS response that we had attributed to RPE in Fig. 3B. To check this, we removed the trials that had catch-up saccades and confirmed that the response to RPE remained robust (Fig. S7D).

In summary, when there were no target jumps, but the value of the target changed, the climbing fibers reported that change, indicating that the future reward would be smaller or larger than expected. Thus, climbing fibers reported both the value of the target, and RPEs.

## RPE multiplicatively enhanced the climbing fiber response to endpoint error

We next quantified the response to endpoint errors while we varied the reward context. In a fraction of the forced trials, during the primary saccade we jumped the target to a random location but did not change its value (Fig. 3C). This produced an SPE without an RPE. As a control, we neither jumped the target nor changed its value.

As the primary saccade ended, the climbing fibers reported the direction of the endpoint error, i.e., an increased CS rate when the error was in direction *θ*, and a suppression when it was in direction *θ* + *π* (Fig. 3D). The encoding of endpoint error was similar to but muted as compared to the CS response to the primary target (Fig. S8A): the preferred direction *θ* remained stable (Fig. S8B), and like the response to the primary target, the response to error was multiplicatively affected by value of the secondary target (Fig. 3D).

To quantify the timing of encoding of direction vs. value, we compared the CS rates in the high and low reward conditions when the error was in direction *θ* vs. *θ* + *π*. The results (Fig. 3E) revealed that the direction of the error vector was reported at 63.8±0.6ms following completion of the primary saccade, and value information was reported at 80.9±0.9ms. Thus, reward information arrived 24ms after direction information, consistent with our observations in Fig. 1G.

We next considered forced trials in which both the target jumped, and its value changed, i.e., SPE & RPE (Fig. 3F). The critical question was whether the CS response to the secondary target merely encoded its reward value, or did it also report an RPE. In both the LH and HH conditions there was an endpoint error. However, in the LH condition the value of the trial had increased, whereas in the HH condition it had not. We compared the CS response in the LH condition with the HH condition and found that the CS rates dramatically increased in the LH condition, but this increase was smaller in the HH condition (Fig. 3F, LH vs. HH, t(903)=8.86, p=4.3E-18).

As a result, the encoding of error direction was multiplicatively modulated if the secondary target signaled high reward (HH condition), and modulated more if that error was in the RPE+ context (LH condition, Fig. 3F, right plot). In contrast, if the reward was less than expected (HL condition) the resulting RPE-reduced the CS response as compared to the LL context (Fig. 3F, HL vs. LL).

Finally, we considered the possibility that the climbing fibers reported not only the direction of the error vector, but also its amplitude ^38^. We projected the endpoint error vector onto direction *θ* and found that the CS rates progressively grew as this projection increased from negative to positive values (Fig. 3G left subplot). Notably, the reward context multiplicatively modulated this response, producing the largest CS rates in the RPE+ condition (Fig. 3G, right subplot).

Thus, the climbing fibers reported both the direction and magnitude of the endpoint error. The directional encoding was multiplicatively enhanced for high valued targets and enhanced further in the presence of an RPE+ event.

## The P-cell’s potent vector was aligned with the preferred direction of its climbing fiber

We can summarize the climbing fiber response to stimulus reward value *r*, RPE *δ*, and endpoint error ***e***, with the following equation:

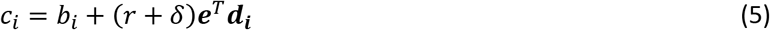

In this equation, *c_i_* is the CS firing rate in P-cell *i*, and *b_i_* is its baseline value. This equation explains that the climbing fiber input exhibits a directional tuning because, analogous to gradient descent, it is transmitting the dot product of the reward-dependent error vector ***e*** upon the P-cell’s potent vector ***d_i_***. If this is true, then ***d_i_*** should be a vector that points in the preferred direction of the climbing fiber, i.e., *θ_i_*. Moreover, as the magnitude |***d****_i_*| increases, i.e., the P-cell has a greater ability to displace the eyes, so should the depth of modulation of the CS tuning function. We set out to test these predictions by measuring ***d_i_***.

To estimate the potent vector ***d_i_*** for each P-cell, we relied on the fact that CSs were low frequency (1Hz), stochastic events that completely suppressed the P-cell’s ability to generate simple spikes (Fig. 4A, left subplot). When the CS event occurred during primary saccades (probability 0.079±0.0012), the resulting SS suppression was not different for high or low valued target (SS rate, change from [-10 to −1]ms before CS to [+1 to 10]ms after CS, pairwise t-test, p=0.36). However, the suppression produced a slight change in eye trajectory, acting as a stochastic perturbation ^22,39,40^. We performed spike-triggered averaging, comparing saccades that had the same starting position and were made toward the same target, but did or did not experience a suppression (Fig. 4B). The difference in the eye trajectories in velocity space, i.e., Δ***ẏ***(*t*), was a vector that remained parallel to *θ_i_* throughout the duration of the saccade (Fig. 4B). When two P-cells with similar climbing fiber preferred directions, i.e., 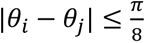, were suppressed during the same saccade, the amplitude of the disturbance more than doubled, but once again Δ***ẏ***(*t*) remained parallel to *θ*. Critically, these effects on saccades were not different for the high or low valued targets (velocity change at peak speed, high value 3.61±0.39 deg/s; low value: 3.17±0.42 deg/s; mean±SEM, pairwise t-test, p=0.42), demonstrating that the potent vector ***d****_i_* remained constant despite changes in the reward context. We next compared the preferred direction *θ_i_* and the direction of its potent vector ***d****_i_* (Fig. 4C). The angular difference exhibited a unimodal distribution with a mean value near zero (6.52±3.3 deg, mean±SEM).

**Fig. 4.**
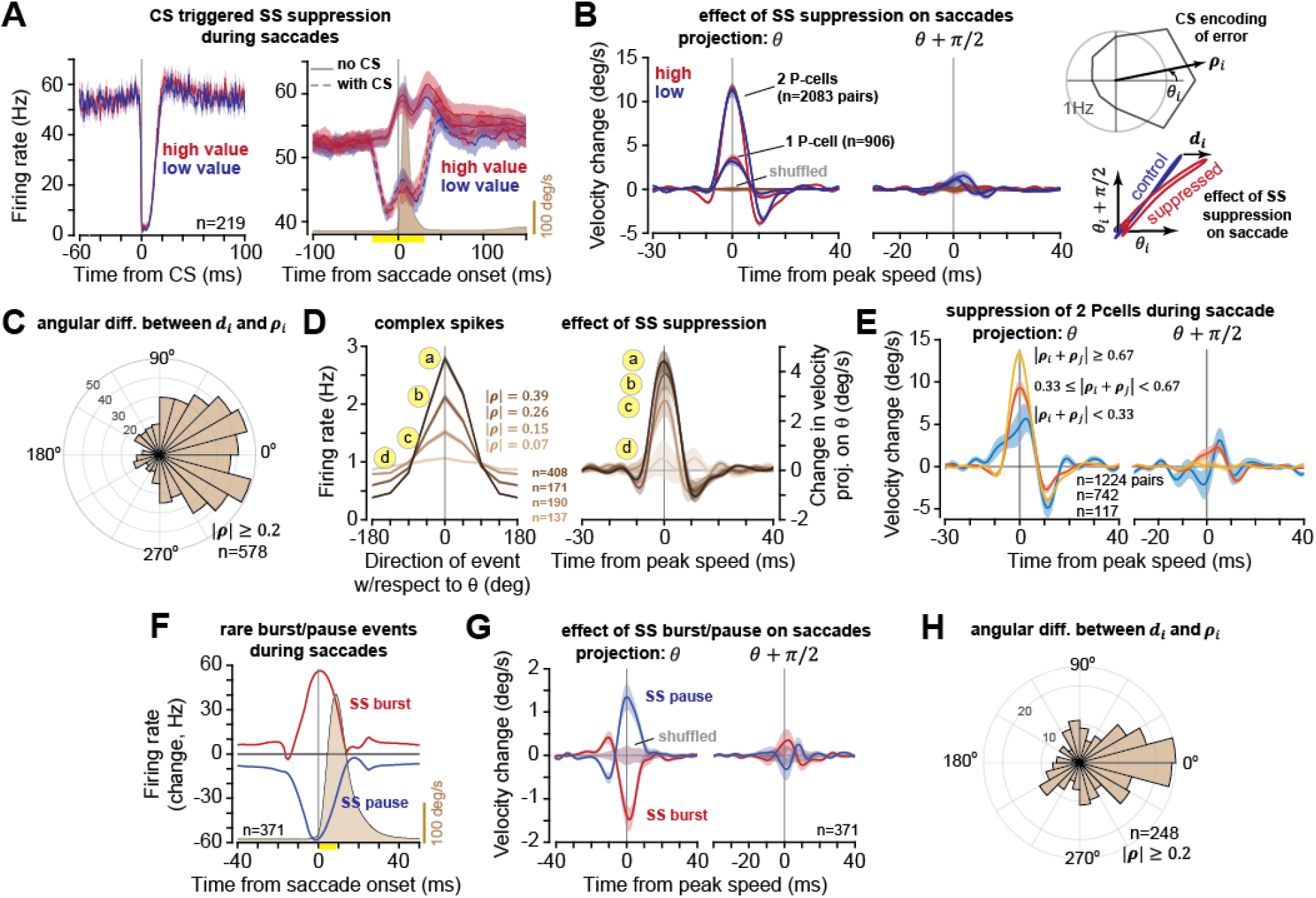
The P-cell’s potent vector was aligned with the preferred direction of its climbing fiber. **A**. Left: simple spike (SS) firing rates aligned to CS onset during saccades. Right: SS rates during saccades that did or did not experience a CS. **B**. Suppression induced change in saccade velocity parallel or perpendicular to the preferred direction of the climbing fiber input, i.e., *θ*. The change in velocity is plotted for suppression of one P-cell, and for simultaneous suppression in two P-cells in which 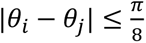. The suppression induced trajectory change ***d****_i_* was computed at peak saccade speed. ***d****_i_* is labeled as the potent vector of P-cell *i*. **C**. Distribution of angular difference between ***d****_i_* and climbing fiber tuning vector ***ρ****_i_*. **D**. As the depth of modulation in the climbing fiber tuning increased, reflecting an increase in |***ρ***|, so did the amplitude of the suppression-induced displacement |***d***|. **E**. Effect of simultaneous suppression of two P-cells, conditioned on |***ρ****_i_* + ***ρ****_j_*|. The suppression induced displacement tended to be larger in P-cell pairs where the climbing fiber inputs had a larger ***ρ*. F.** We identified saccades in which a P-cell produced statistically rare simple spike bursting or pausing during 10ms onset period. This plot shows change in SS rates, with respect to control trials, for these saccades. **G**. Burst or pause induced change in saccade trajectory. **H**. Direction of the potent vector as estimated via the burst/pause analysis vs. *θ*. Error bars are SEM, except as noted for 95% confidence intervals.

In summary, when P-cell *i* was briefly suppressed during a saccade, the result was a displacement of the eyes along a vector ***d****_i_*. This displacement was on average parallel to the preferred direction *θ_i_* of its climbing fiber, with an amplitude that was invariant to the reward context.

## The amplitude of the potent vector was correlated with the depth of modulation of the complex spikes

We represented the directional encoding in each climbing fiber as a vector by fitting a von Mises distribution to the CS rates (Fig. 4B, right top plot), resulting in a vector ***ρ****_i_* with direction *θ_i_* and magnitude |***ρ****_i_*|. Eq. (3) predicted that the depth of the CS tuning, i.e., |***ρ****_i_*|, would be proportional to the amplitude of the potent vector |***d****_i_*|. That is, P-cells that had a strong influence on eye movements, as reflected in the large |***d_i_***|, should also have a climbing fiber input that was strongly modulated as a function of the direction of the error. We binned the climbing fibers based on the magnitude |***ρ****_i_*| into four groups (Fig. 4D) and found that as |***ρ****_i_*| increased, so did the displacement that was produced by the SS suppression (Fig. 4D, right subplot, ANOVA F(3)=7.23, p = 8.4E-5). Moreover, when two P-cells were simultaneously suppressed during a saccade, with 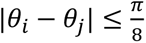, the resulting change in saccade velocity tended to grow with the amplitude of the sum of the two potent vectors |***ρ****_i_* + ***ρ****_j_*| (Fig. 4E, ANOVA, F(2)=15.72, p=1.68E-7). Thus, the depth of tuning in the climbing fiber that innervated a P-cell grew with the size of its potent vector.

## Control studies

Because the climbing fiber input to a P-cell is the branch of an axon that also makes synapses onto nucleus neurons, we sought independent verification of our estimated potent vectors without reliance on CS-induced P-cell suppression. We looked for statistically rare temporal patterns in each P-cell’s SSs during saccades, i.e., rare pauses or rare bursts that were not associated with a CS, and treated these endogenous fluctuations as perturbations of the cell’s output (Fig. 4F). An SS pause during a saccade displaced the eyes along a vector that had a larger positive component in direction *θ* than direction *θ* + *π*/2 (Fig. 4G, velocity change at peak speed, 1.354±0.298 vs. −0.29±0.26 deg/s; mean±SEM, paired t-test, p=6.97E-6). An SS burst produced a displacement with a larger negative component in direction *θ* as compared to direction *θ* + *π*/2 (Fig. 4G, velocity change at peak speed, −1.383±0.288 vs. 0.31±0.28 deg/s; mean±SEM, paired t-test, p=2.66E-5). Thus, decreases and increases in the same P-cell’s SS activity altered behavior in opposite directions along direction *θ*, confirming that the potent vector of a P-cell was on average aligned with the preferred direction of its climbing fiber (Fig. 4H, angular difference −6.16±5.83 deg), suggesting that statistically rare fluctuations in neural activity may be used to estimate the behavioral effects of individual neurons without externally perturbing them.

## Discussion

We discovered that the information transmitted by climbing fibers combined the subjective, reward-dependent context in which the error was experienced, termed the loss function, with the objective, individual role that the P-cells played in producing the movement, termed the neuron’s potent vector. This suggests that climbing fibers solve the credit assignment problem by providing the gradient of a loss function with respect to the output of the P-cells that they innervate.

We manipulated the loss function by associating reward with visual targets. The climbing fibers responded to target presentation by multiplicatively reporting its direction and reward value via a tuning function. When a saccade ended in error, the climbing fibers reported the spatial properties of the error, as well as its reward context (including RPEs). We used spike-triggered suppression of P-cells to measure each neuron’s downstream effects on saccades and found that the resulting potent vector was colinear with the preferred direction of its climbing fiber. Across P-cells, an increase in the amplitude of the potent vector coincided with an increase in the depth of modulation of the climbing fiber tuning. This implied that the olivary neurons were computing a signal that was proportional to the dot product of the reward dependent error vector and the potent vector of the P-cell that they projected to.

### Climbing fibers combined SPEs with RPEs

The canonical division of labor in computational neuroscience assigns reinforcement learning to the basal ganglia, where midbrain dopamine neurons signal RPEs ^41^, and supervised learning to the cerebellum, where climbing fibers signal SPEs ^42^. Yet, as the brain learns the predictive value of a cue, climbing fibers respond to that cue ^43–45^, especially if the cue predicts future reward ^18^. Indeed, climbing fibers provide a signal that resembles RPEs: they respond to unexpected reward ^20,43^, transfer their response to reward-predicting cues ^19,43^, become less responsive to expected reward ^19,20^, signal omission of reward ^20^ or punishment ^45^, and respond to excitation of dopamine neurons ^46^. These results suggest that single climbing fibers might convey both reward information and sensory prediction errors.

Here, we found that climbing fibers initially reported the spatial information regarding the random visual event, then 25ms later reported its reward value. The timing of these responses resembled the neurons in the superior colliculus^47^, a region that projects to the contralateral inferior olive without maintaining a retinotopic map ^48^. In addition, mesodiencephalic junction (MDJ) receives projections from the frontal and parietal cortex and projects densely to the inferior olive ^49^. Thus, it seems likely that inferior olive neurons are recipient of sensory and reward information from at least the superior colliculus and MDJ. But do climbing fibers encode SPEs, or simply sensorimotor events?

Motor learning studies do not provide a conclusive answer because even if the climbing fiber response declines with repetition of a constant error, that could reflect reduced subjective value (animal becoming sated, Fig. 2B), and not a smaller SPE. However, in classical conditioning, climbing fibers stop responding to an air puff when that event becomes expected ^45^. Dring saccades, climbing fibers respond to endpoint errors, even if those errors do not produce a corrective movement ^29^, and then drive trial-to-trial changes in P-cell simple spikes that accompany changes in saccade trajectory ^14^. These results suggest an encoding of SPEs. But what might be the purpose of combining SPEs with reward and RPEs?

In the context of motor learning, a loss function is a value-dependent, subjective measure of an SPE ^1^. To minimize loss, neurons require a teacher that instructions them on how to change their output, i.e., the gradient of the loss with respect to their output. Here, we found that when a movement ended in error, the climbing fibers multiplicatively combined the spatial properties of that error with its reward context, producing a signal that met the requirements of gradient descent. Thus, our results suggest that the inferior olive receives reward information, and the cues that predict it, because these are modulators of a loss function that instruct the cerebellum regarding how much to learn from error. If so, this could point to use of RPEs for rehabilitation of cerebellar patients ^50,51^.

### How does a climbing fiber know about the P-cell’s potent vector?

Simulations suggest that in a shallow neural network, random weights assigned to the error vector can be an effective teaching signal to neurons in an intermediate layer, given that the downstream weights of those neurons are modified using gradient descent ^52^. The implication of this feedback alignment algorithm is that the olivary neurons may have a random, fixed sampling of the error space, but learning can alter the downstream connections of a P-cell so that those weights match the information that is conveyed in the climbing fiber input. This theory predicts that it is not the climbing fibers that “know” about each P-cell’s potent vector, but rather that the potent vectors gradually adapt to match the preferences of the climbing fibers.

There are two kinds of evidence for this conjecture. Genetic re-routing of climbing fiber input during development leads to reorganization of P-cell and molecular layer interneuron activities, without changes in the mossy fiber inputs ^53^. In the developing zebra fish, changing the activity of inferior olive neurons changes the downstream connections of P-cells ^54^. These results suggest that climbing fibers serve as teachers for two different circuits: the parallel fiber input to P-cells, and the downstream network that connects the P-cells to behavior ^54^. In this framework, the climbing fiber response to unexpected sensory events guides training of parallel fiber synapses, but the climbing fiber activity during self-generated movements trains the pathways that connect the P-cells to behavior, possibly through the effects that synchronized suppression of P-cells have on nucleus neurons ^55^.

### Limitations and predictions of the theory

Our theory predicts a limitation of cerebellar learning: because climbing fibers encode *dL*/*dp_i_* and not *dL*/*dw_ji_*, they cannot differentially guide learning in the parallel fiber synapses that happen to be active at the same time. This means that during a saccade, as the mossy fibers that encode goal of the movement are co-active with the ones that encode the motor commands ^23^, endpoint error will produce learning in both types of granule cell inputs, conflating errors in motor commands with errors in representation of goals.

Climbing fibers should be active in response to movements that result in error. We induced errors by jumping the target after primary saccades, and there was indeed robust CS modulation in response to the jumped target. However, CS modulation was equally strong in response to appearance of the primary target, which was not a movement error. This suggests that the climbing fiber responds to all unexpected visual events, i.e., SPEs. Because CSs can only induce plasticity at recently active parallel fiber synapses ^31^, when a movement ends in error, the resulting CSs induce plasticity at parallel fiber inputs that were active during the preceding movement. However, CSs that occur in response to primary target are likely to induce no plasticity, as the relevant parallel fibers were not active during fixation. Thus, climbing fibers can respond to SPEs, but learning will still only occur when an unexpected event was the result of a movement error. Additionally, as mentioned above, the CSs that result from primary saccades may serve to instruct downstream plasticity to keep the potent vector aligned with the climbing fiber’s preferred direction.

In trials in which we changed the target value, but not its position (Fig. 3A), climbing fibers still responded to the RPE (Fig. 3B). Because saccades are generally hypometric, even without a target jump there are endpoint errors. The fact the CS response to target value change was larger when the primary saccade was in direction *θ*, and thus the endpoint error was also in direction *θ*, appears consistent with this possibility.

Finally, the CS response was largest for endpoint errors in the context of RPE+ and smallest in the context of RPE- (Fig. 3F). This matched behavioral measures of trial-to-trial learning (Fig. 1D). In contrast, the CS response was also larger for high valued targets with respect to low valued targets, but our behavioral measures found no difference in these two contexts. Other experiments in which target directions repeat have found that high reward does improve rates of learning ^4^. Because our experiment relied on random presentation of target directions, this limited our ability to measure trial-to-trial change. It is possible that learning from error may rely on multiple regions of the cerebellum, and climbing fiber activity in the vermis may not be a perfect predictor of change in behavior.

## Expanding the current theories of biological learning

Our results suggest that the cerebellum solves the credit assignment problem in a way that differs from the computational frameworks currently proposed for biological learning. In the Marr–Albus theory, climbing fibers carry a global teaching signal that broadcasts errors to P-cells ^56,57^, thereby inducing synaptic plasticity in cells that happen to be active at the time of the error. However, P-cells are generally active across all directions of a given movement (e.g., saccades ^32^, licking ^58^, or reaching ^59^), making it unclear how a common error signal is transformed into a neuron-specific teaching signal.

A second class of theories, inspired by reinforcement learning, avoids explicit error assignment by treating neural variability as stochastic exploration, modifying synapses according to correlations between perturbations and changes in reward ^60,61^. While such policy-gradient mechanisms can learn without detailed knowledge of downstream circuitry, and have been proposed for the cerebellum ^62^, they are inefficient because they estimate gradients through trial-and-error sampling ^10^.

In contrast, artificial neural networks solve credit assignment through backpropagation ^9^, where every neuron receives the gradient of the loss function with respect to its own output. Our results suggest that the cerebellum approximates this latter computation without implementing backpropagation itself.

Thus, the inferior olive provides a surprisingly sophisticated solution to the credit assignment problem. The signal in the climbing fiber is not merely a prediction error, nor merely a reward signal, but a neuron-specific gradient that instructs the P-cell on how to change its activity to reduce future loss.

## Supplementary Materials

### Experimental Methods

We collected data from two marmosets during a 2.5 year period, *Callithrix Jacchus*, 350-370 g, subjects 132F (M1, Charlie, Male, 7 years old), and 65F (M2, Barney, Male, 7 years old). The marmosets were born and raised in a colony that Prof. Xiaoqin Wang has maintained at the Johns Hopkins School of Medicine since 1996. Our procedures were approved by the Johns Hopkins University Animal Care and Use Committee in compliance with the guidelines of the United States National Institutes of Health.

#### Data acquisition

Following recovery from head-post implantation surgery ^63^, the animals were trained to make saccades to visual targets and rewarded with a mixture of applesauce and lab diet ^64^. They were trained on 4 visual targets, each an abstract shape (Fig. 1C): two were associated with relatively large amounts of applesauce, and two were associated with smaller amounts. These visual targets were presented on an LCD screen with 500Hz refresh rate and low latency (Dell AW2524H). Binocular eye movements were tracked at 1000 Hz using the EyeLink system.

We performed MRI and CT imaging on each animal and used this data to design an alignment system that defined trajectories from the burr hole to various locations in the cerebellar vermis ^63^, including points in lobule VI and VII. We used 3D Slicer software to align the T2 MRI to the marmoset atlas ^65^. We then aligned the CT to the transferred T2 MRI. We used a piezoelectric, high precision microdrive (0.5 micron resolution) with an integrated absolute encoder (M3-LA-3.4-15 Linear smart stage, New Scale Technologies) to advance the electrode. To reach the cerebellar cortex, we planned a posterior burr hole and avoided the confluence of sinuses using MRI T2 images. This approach enabled us to record from multiple folia in lobules VI and VII simultaneously.

We recorded from the cerebellum using Neuropixels 1.0 NHP probes as well as 64-channel checkerboard or linear high-density silicon probes (M1 and M2 probes, Cambridge Neurotech). For Neuropixels we used the National Instrument acquisition system (NI PXIe-1071). Data were sampled at 30 kHz and aligned to the eye tracking system time as the reference time using random TTL signals. For the Cambridge probes we used a 64-channel head stage amplifier and digitizer (RHD2132 and RHD2164, Intan Technologies, USA), then connected the head stage to a communication system (RHD2000 Evaluation Board, Intan Technologies, USA). Because the conductive coating on the Cambridge probes degraded after each insertion into the brain, we re-coated the probes and restored their low impedance after every 3-4 recording sessions ^66^. For accurate reaction time and visual target time measurements, we measured the video monitor latency in real-time with a photodiode.

#### Experiment design

We employed two types of trials: forced trials (80%), and choice trials (20%) (Fig. S1). Forced trials measured the neural and behavioral responses to sensory and reward prediction errors. Choice trials measured the subjective value that the marmosets had learned to assign to each target. Choice and forced trials were randomly intermixed.

##### Forced trials

(Fig. S1A). These trials began with fixation of a center target (a gray square). Next, a primary target (one of the 4 abstract images) appeared at one of 8 randomly selected directions at a displacement of 4-5 deg. Each visual target was 2deg in diameter. We had four types of forced trials: 1) target jump only, 2) target jump & value change, 3) value change only, and 4) no target jump & no value change.

#### Target jump only trials

(Fig. S1C, 30-36% of forced trials). As the subject made a saccade to the primary target, that target was erased and a secondary target was presented at a distance of 2.5-3.5 deg at one of 8 randomly selected directions. The displacement of the target mid-saccade was a sensory prediction error (SPE) in the sense the target was not on the fovea when the primary saccade concluded. We label this event as an SPE because it produced error-dependent learning, as evidence by a change in the trajectory of the subsequent primary saccade (Fig. 1D) ^14^. In these trials the primary and secondary targets were visually identical. Following the completion of the secondary saccade, a tone sounded to indicate the amount of food earned (high or low), and then the food was dispensed.

##### Target jump & value change trials

(Fig. S1C, 20-22% of the forced trials). In these trials, the secondary target had a different value than the primary target. For example, if the primary target was of high value, but the secondary target was of low value, the successful completion earned only a small amount of food. This, we hypothesized, produced a negative RPE. On the other hand, if the primary target was low value, but the secondary target was high value, successful completion now earned a large amount of food, producing what we hypothesized was a positive RPE.

##### Value change only trials

(Fig. S1C, 20-22% of the forced trials). As the primary saccade took place, we changed the target image but not its location. Thus, the target might switch from high to low, resulting in RPE-, or from low to high, resulting in RPE+.

##### No target jump & no value change trials

(Fig. S1C, 30-36% of the forced trials). These were control trials in which both the location and value of the primary target remained unchanged.

##### Choice trials

(Fig. S1B). These trials were randomly intermixed with force trials. A choice trial began with fixation of the center target. Two targets were displayed, one with high value and the other with low value. The locations of these targets were opposite of each other (Fig. S1B). The two targets were placed along one of 4 randomly chosen axes with respect to the center target. The subjects indicated their choice via a saccade and were then rewarded by the amount specified by the chosen target. Thus, the choices that the subjects made in these trials allowed us to quantify the relative value that they had assigned to each target.

Reward was dispensed following successful completion of each trial. However, because the food increment was small, the subjects chose to work for a few consecutive trials, tracking the visual targets and allowing the food to accumulate, then stopped doing the task and harvested the food via a bout of licking ^64^.

#### Visual targets

We used four abstract shapes (2 deg in diameter) as visual targets for the primary and secondary saccades (Fig. 1C): two of the targets were associated with high reward, while the other two were associated with low reward. We balanced the stimuli in the two monkeys so that the two high valued stimuli in one monkey were low valued stimuli in the other monkey. Monkey M1 was presented with only 2 stimuli in each session, while monkey M2 was presented with all 4 stimuli in each session.

#### Identification of saccades

We analyzed eye movements using a deep neural network that detected both saccades and microsaccades ^67^. The pre-trained networks for human and macaque monkeys did not perform well for marmosets. Hence, we designed a custom Matlab GUI-based program to curate the saccades (https://github.com/ShadmehrLCMC/SACCURATE). We then re-trained the network through transfer learning for each individual animal using seven 30-45 minutes of recording. To prevent any potential false positive saccades, we pruned the saccades by fitting a bivariable Gaussian distribution to two different feature spaces defined as biologically relevant metrics ^68^ and removed saccades that were outliers (less than 1% chance of belonging to the distribution). The two feature spaces were the log-log main-sequence plots (maximum velocity vs. amplitude) and the log-log acceleration time to deceleration time ratio vs. amplitude of saccade. We then analyzed and found valid fixations among candidate fixations after or before each detected saccade. We measured and used data-driven thresholds on steady eye position criteria including maximum displacement, dispersion on x and y axes, fixation duration as well as maximum velocity. We discarded fixation candidates with lost eye signals due to instability of eye tracking or blinking.

We detected the onset and offset times of each saccade using a trained neural network, as described above. We then low pass filtered the eye position traces with a 3rd order Butterworth filter with 100 Hz cut-off frequency. We then calculated the saccade velocity by differentiating the eye position trace and found peak velocity times and values.

#### Catch-up saccades

In the No Target Jump condition of the forced trials, there was only a primary target without a secondary target. In this case the primary saccade was occasionally followed by a small “catch-up” saccade. We labeled catch-up saccades based on the following criteria. 1) The onset of the candidate saccade had to occur within 0.5° of visual angle from the endpoint of the preceding saccade, ensuring that it was part of the same movement sequence. 2) The amplitude of the candidate saccade had to be less than 2.5° of visual angle, consistent with a corrective adjustment rather than a new primary movement. 3) The movement vector of the candidate saccade had to fall within 30° of the vector pointing from its start position toward the primary target. 4) The candidate saccade was required to reduce the distance between gaze position and the primary target. Specifically, the Euclidean distance to the target at saccade offset had to be smaller than the distance at saccade onset. 5) The latency between the offset of the primary saccade and the onset of the candidate saccade had to be ≤ 500 ms.

#### Identification of cell types

We used OpenEphys ^69^ for electrophysiology data acquisition, and then used Kilosort 2.0 and Phy 2.0 ^70^ to manually curate the spikes. Each recording was curated twice by two experienced neurophysiologists.

We performed manual cell type identification through identification of the layers of the cerebellar cortex. First, we identified the P-cells via CS-induced SS suppression (Fig. S3A). As P-cells are large cells, their spike waveforms are present on multiple channels of the silicon probes (Fig. S3B, SS near dendrites and soma). Next, using the CS waveforms on the channels we identified the orientation of the P-cell’s axon and dendrites, thus identifying the molecular, Purkinje, and granular layers ^23,55,71,72^.

For example, we identified the molecular layer via the downward, broad CS waveforms in the dendrite tree of the P-cells (Fig. S2B, complex spikes near dendrites). We controlled for contamination in the cross-correlograms in P-cell simple and complex spikes and removed the spikelets and double detection of CSs as SSs by aligning the two waveforms to each other and removing the copies in SS units

#### Quantifying the quality of the neuronal recording

A summary of the measures used to verify the quality of the neurophysiological data is provided in Fig. S3A. To measure the isolation quality of each neuron, we computed the conditional probability Pr(*s*(*t* + Δ) = 1|*s*(*t*) = 1), that is, the probability of a spike at time delay Δ, given that the neuron produced a spike at time zero. We then multiplied this probability by 1000 (bin size 1 ms) and plotted the results as firing rate. For well-isolated cells, we expected a clean refractory period. To measure the noise rate in each neuron’s spiking, we quantified this conditional probability at Δ = 1ms, except for complex spikes, for which we used a 5ms period. For example, to quantify the quality of the P-cells, we measured the noise rate and found that for the simple spikes, this rate was 0.63±0.47 Hz (median ± median absolute deviation, MAD). For the complex spikes, the noise rate was 0±0 Hz.

#### Measuring the climbing fiber encoding of sensorimotor events

In lobules VI and VII of the cerebellar vermis, the climbing fibers report the direction of visuomotor events ^73,74^, likely because of the superior colliculus projections to the inferior olive ^48^. For example, the climbing fibers reported to the cerebellum both the direction of the visual event, and then independently, the direction of the planned saccade ^29^. We used the properties of the climbing fiber input to quantify the CS tuning (Fig. S4). The procedures were identical to those described in ^23^. Briefly, we quantified this encoding by measuring the CS rate as a function of the direction of the visuomotor event during two different windows: 40 to 85 ms with respect to the visual cue onset, and −70 to +30 ms window from saccade onset. We then fitted a Von Mises distribution to the resulting rates as a function of angle ^75^. This produced two vectors, ***ρ****_v_* that quantified the strength and direction of the CS response to the visual events, and ***ρ****_m_* that quantified the strength and direction of the CS response to the motor events. Each vector had an angle *θ* and amplitude |***ρ***|, where the amplitude identified the sharpness of tuning. Thus, the amplitude of the vector was bounded by 0 (not tuned), and 1 (maximum sharpness). This produced two distinct vectors for each P-cell: one describing the tuning with respect to visual events, and one describing the tuning with respect to motor events. However, encoding of these two vectors were very similar: the magnitude of the vector for the visual event was highly correlated with the magnitude of the vector for the motor event. Hence, for each P-cell we used the vector with the larger amplitude and labeled it as its CS tuning vector ***ρ***.

#### Computing a potent vector for each P-cell using naturally occurring perturbations

A neuron’s potent vector describes the direction and magnitude of the downstream behavioral effect produced by a change in that neuron’s output. We estimated the potent vectors of P-cells using spike-triggered averaging of naturally occurring perturbations in SS activity during saccades. We first used CS-induced suppression of simple spikes as a physiologically defined perturbation. We then applied a complementary method that identified statistically rare pauses and bursts in SS activity without relying on CSs. These analyses each produced two related but distinct measures for each neuron. First, we calculated a time-resolved velocity perturbation, Δ**v**(*t*), which described how the effect of the neural event evolved throughout the saccade. We then used this quantity to compute a single two-dimensional potent vector ***d*** that summarized the horizontal and vertical velocity perturbation over a predefined post-saccadic interval. For perturbation and control trials, the time-resolved velocity difference was

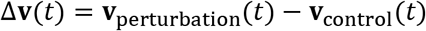

where 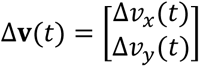. For each target direction *j*, ***d*** was calculated by averaging Δ**v**(*t*) over a brief window after saccade onset (see below). The final potent vector for a P-cell was obtained by averaging ***d****_j_* across the valid target directions.

#### Potent-vector estimation for CS-induced SS suppression in single P-cells

We focused on primary saccades because these movements began near the central fixation point and had a fixed amplitude. Saccades were divided into two groups: (1) saccades in which the P-cell did not experience a CS during the time window −60 to +40ms with respect to saccade onset (NOCS: no cs), (2) saccades in which the P-cell experienced a single CS during the time window −30 to +30ms with respect to saccade onset (WCS: with cs). The wider exclusion interval for no-CS trials prevented complex spikes near the analysis window from contaminating the control group.

The horizontal and vertical velocity traces were first adjusted for initial eye position by regression (see below). We then calculated the CS-induced velocity perturbation:

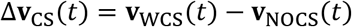

For the time-series analysis, Δ**v**_CS_(*t*) was projected onto the preferred direction of the climbing-fiber response, *θ*, and onto the orthogonal direction, 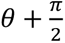. The projected velocity differences were first computed separately for each saccade direction and were then averaged across directions. For potent-vector estimation, the Cartesian velocity difference was averaged over 0–40 ms after saccade onset for each saccade direction. These vectors were subsequently averaged across valid directions to produce the CS-derived potent vector ***d****_CS_*. To determine whether the CS-derived potent vector was aligned with the climbing fiber’s preferred direction *θ*, we calculated the wrapped angular difference.

#### Potent-vector estimation from CS-induced SS suppression in pairs of P-cells

For simultaneously recorded P-cell pairs, each cell had a climbing-fiber tuning vector, 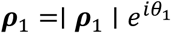 and 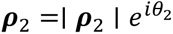, where *θ*_1_ and *θ*_2_ were the climbing-fiber preferred directions and the magnitudes represented the depths of directional tuning. Cell 1 was defined as the cell with the larger tuning magnitude. Pairs were included when 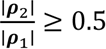. This criterion prevented the contribution of the weakly tuned cell from being negligible relative to that of the more strongly tuned cell. The angular separation between *θ*_1_ and *θ*_2_ was required to be no greater than *π*/8. We then divided the saccades that were made from the same initial position to the same target into two groups: (1) saccades during which neither of the two P-cells experienced a CS (NOCS, −60 to +40ms window with respect to saccade onset), (2) saccade during which both of the P-cells experienced a single CS within 30ms of each other (WCS, −30 to 30ms window). After regression adjustment for initial eye position, the pairwise velocity perturbation was:

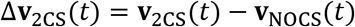

This velocity difference was projected onto *θ*_net_ and *θ*_net_ + *π*/2. For analyses of perturbation magnitude, pairs were grouped according to the magnitude of the net CS-tuning vector, ∣ ***ρ***_net_ ∣.

#### Identification of rare pause or burst events

To estimate potent vectors without relying on CSs, we developed an analysis based on trial-to-trial variability in P-cell SS output. The method was motivated by the observation that CS-induced SS suppression can be viewed as a rare pause event: during the suppression, the P-cells temporarily fail to produce their expected output, resulting a small but measurable change in the movement.

For each trial, we calculated an age function, *a_i_*(*t*), which represented the elapsed time since the most recent simple spike at each time *t*, i.e., a function that grew with the inter-spike interval. The pausiness score *P_i_* for trial *i* was defined as

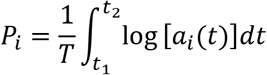

where *t*_1_ and *t*_2_ defined the detection window and *T* = *t*_2_ − *t*_1_. The log function imposed diminishing returns as the inter-spike interval increased, reducing the disproportionate influence of exceptionally long pauses. The pausiness score quantified how sparse and temporally separated the simple spikes were during the detection window. The detection window extended from 0 to 10 ms after saccade onset. Because SS activity depended on movement direction and target reward context, rare events were identified separately within each direction and context. Trials in the upper 20% of the pausiness-score distribution were classified as rare pause events, whereas trials in the lower 20% were classified as rare burst events. Trials containing a CS within the CS exclusion interval were excluded from this analysis. Thus, rare events represented fluctuations in SS activity that occurred independently of CS-induced suppression.

After classifying trials, we used the same logic used for CS-induced suppression. Horizontal and vertical velocity traces were first adjusted for initial eye position using regression. The velocity perturbation associated with a rare pause was

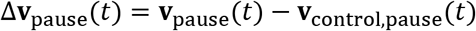

whereas the perturbation associated with a rare burst was

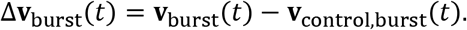

For the time-series analyses, both velocity differences were projected onto the climbing-fiber preferred direction *θ* and the orthogonal direction, *θ* + *π*/2. A rare pause represented a transient reduction in P-cell output and was therefore expected to produce a perturbation with the same direction as CS-induced suppression. A rare burst represented a transient increase in P-cell output and was expected to produce an oppositely directed perturbation.

#### Potent-vector estimation using rare pauses and bursts

For each target direction *j*, a pause-derived potent vector was calculated by averaging the Cartesian pause-associated velocity change from 0 to 25 ms after saccade onset:

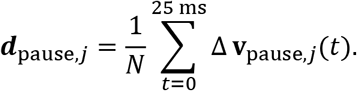

A burst-derived potent vector was calculated in the same manner. Because the burst-associated behavioral effect was expected to have the opposite sign from the effect of reducing P-cell output, the burst-derived vector was sign-reversed. The pause-derived vector and sign-reversed burst-derived vector were then averaged within each direction and then were averaged across valid saccade directions. To determine whether the rare-event method recovered the same potent vector direction as the CS-induced suppression method, we calculated the wrapped angular difference.

#### Datasets used for perturbation analysis

The CS-induced SS suppression analyses were performed using the data collected in the current study. For the rare spike event analysis, to verify its validity we analyzed both the data collected in the current study, and the data that we had previously recorded ^23^. In the analysis of the current data set, because the primary target had two reward conditions, we analyzed each reward condition separately and then averaged the results across the reward conditions.

#### Regression adjustment for initial eye position

Small differences in initial eye position could produce systematic differences in the subsequent saccade trajectory. We removed the small trajectory biases caused by differences in the initial eye position before comparing perturbation and control trials. This regression was performed separately for each P-cell, target reward condition, and saccade direction. At each time point, horizontal and vertical eye velocity were modeled as linear functions of the horizontal and vertical initial eye position:

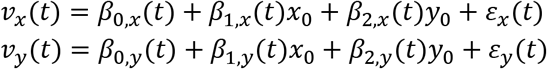

where *x*_0_ and *y*_0_ are the horizontal and vertical eye positions at the beginning of the saccade. The residual velocity traces, *ε_x_*(*t*) and *ε_y_*(*t*), were used in all subsequent perturbation analyses.

#### Computing latency to encode direction and value of the visual stimulus

The climbing fibers reported both the direction and value of the visual stimuli. To compute the timing of these two pieces of information, we used a bootstrap procedure. For each bootstrap iteration, P-cells were resampled with replacement from the full population, and the population-average response trace was recomputed. This was done separately for the direction encoding, defined as *θ* − (*θ* + *π*), and the value encoding, defined as high-reward minus low-reward trials in direction *θ*. For each resampled population trace, baseline activity was measured from −50 to 0 ms relative to target onset. A detection threshold was defined as the baseline mean plus three times the baseline standard deviation. Latency was defined as the first post-target time point at which the response exceeded this threshold for at least 5 consecutive milliseconds. This procedure was repeated 100 times, producing a bootstrap distribution of latency estimates for both direction and value signals. The reported latency is the mean of the valid bootstrap latencies, and the uncertainty is the SEM across valid bootstrap samples.

#### Statistical testing

We performed t-tests, or rank sum tests, or ANOVAs, to compare distributions. For population velocity traces, null distributions were generated by randomly permuting perturbation and control labels within each P-cell or P-cell pair, target reward condition, and saccade direction. The number of trials assigned to the perturbation and control groups was preserved. The complete averaging procedure was repeated for each permutation, including averaging across target reward conditions, directions, and cells or pairs. This procedure was repeated 1,000 times. Pointwise confidence intervals were obtained from the resulting null distributions. Observed population traces were computed using the same averaging hierarchy as the permuted traces. Unless otherwise indicated, velocity differences were first calculated within target condition and saccade direction, then averaged across target conditions within direction, across directions within each cell or pair, and finally across cells or pairs.

---

**Fig. S1.**
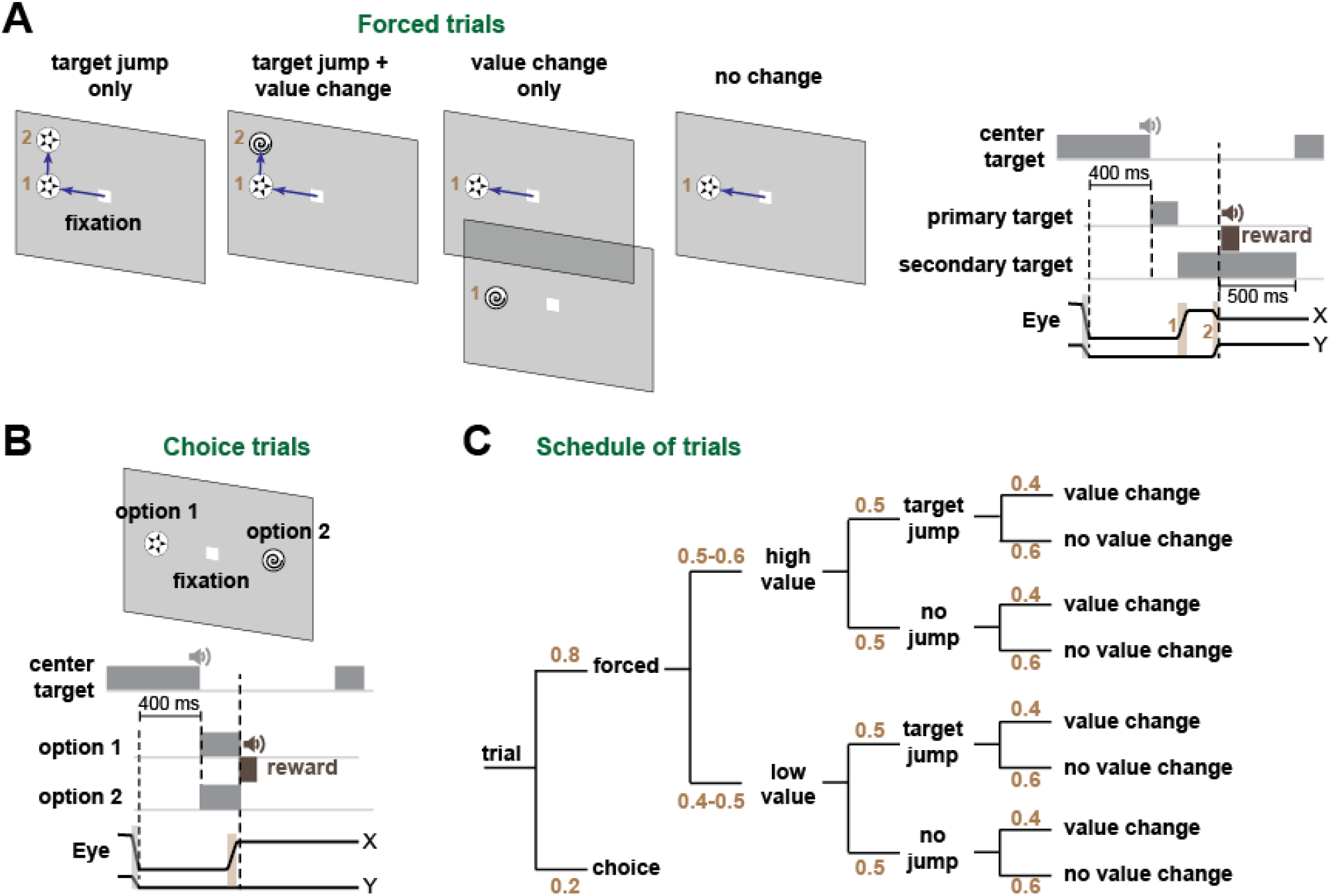
Experiment design. **A**. There were four types of forced trials: 1) Trials with target jump only (primary target followed by a secondary target without a change in target value). This produced an endpoint error without an RPE. 2) Trials with target jump and value change. This produced an endpoint error with RPE. 3) Trials with value change only. This produced an RPE without an endpoint error. 4) Trials without target change. In target jump trials, the secondary target was presented at the detection onset of the primary saccade. **B.** Choice trials. **C**. Schedule of trials. The numbers indicate probability.

**Fig. S2.**
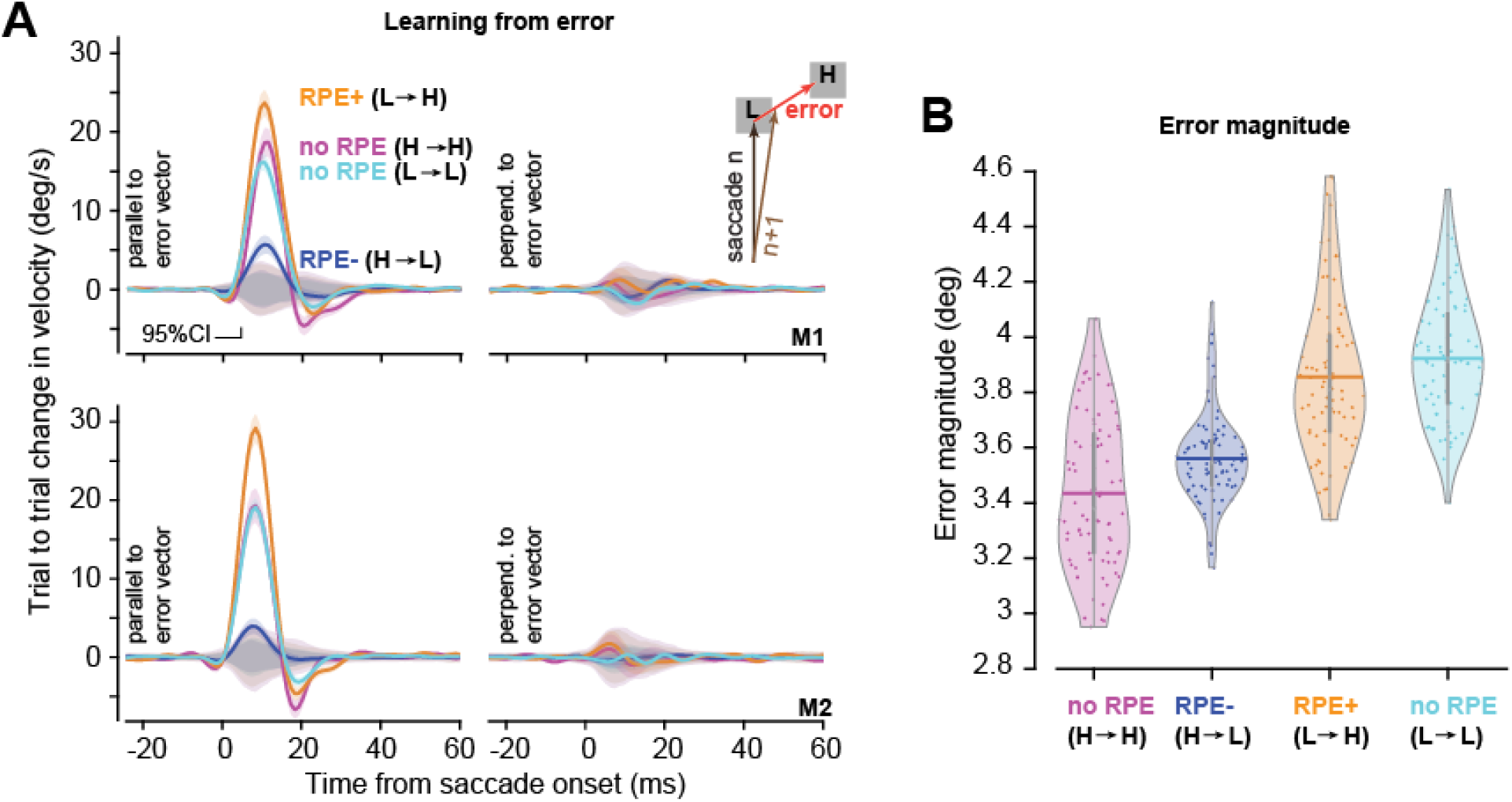
Learning from saccade endpoint error in each subject. A. In target jump trials, subjects made a primary saccade and experienced an endpoint error, defined as the position of the secondary target with respect to the primary saccade endpoint. The plots show trial-by-trial change in velocity of primary saccades, projected onto a unit vector parallel to, or perpendicular to the error vector. The learning was largest in the LH condition (RPE+), smallest in the HL condition (RPE-). **B**. Error magnitude as a function of trial type. Each dot is a single session.

**Fig. S3.**
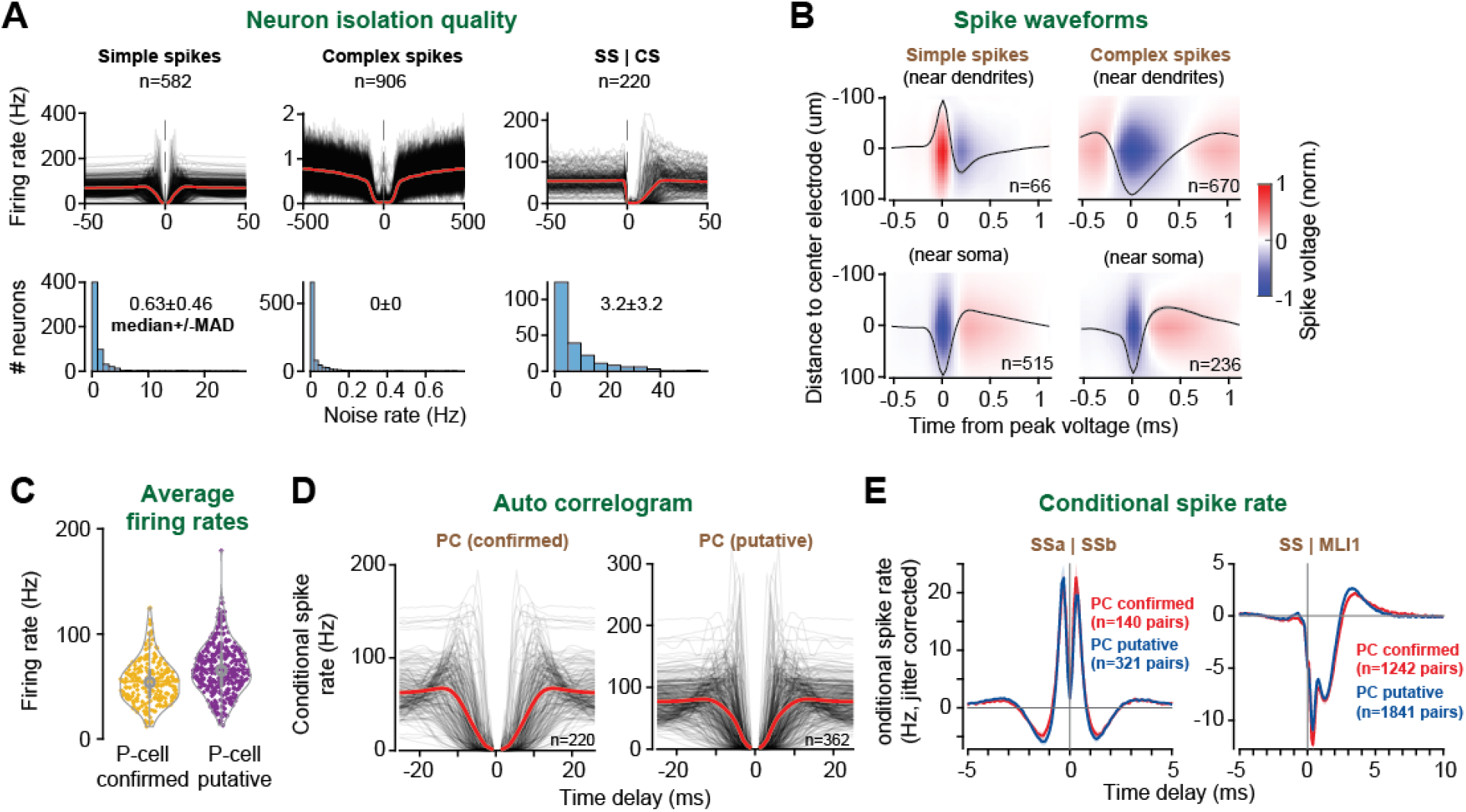
Isolation quality of neurons in the database. **A**. Top row: autocorrelation of simple spikes, complex spikes, and conditional spike rate of simple spikes given a complex spike at time 0. Bottom row: noise rate in each neuron’s spiking, quantified via violation of refractory period, measured via conditional probability at time delay Δ = 1ms, except for complex spikes, for which we used a 5ms period. **B**. Waveforms for simple and complex spikes near the P-cell’s soma and dendrites as a function of their distance to the center electrode. The black trace indicates the spike shape at the center electrode. The upward spike shape for simple spikes, and slow downward shape for the complex spikes, identify the P-cell’s dendrites and therefore the molecular layer. **C**. Average firing rates of confirmed and putative P-cells. **D**. Auto correlogram of confirmed and putative P-cells. **E**. Left: average conditional spike rate for pairs of confirmed and putative P-cells. The high rates of synchrony are present in both groups. Right: MLI1 inhibition of confirmed and putative P-cells. Error bars are SEM.

**Fig. S4.**
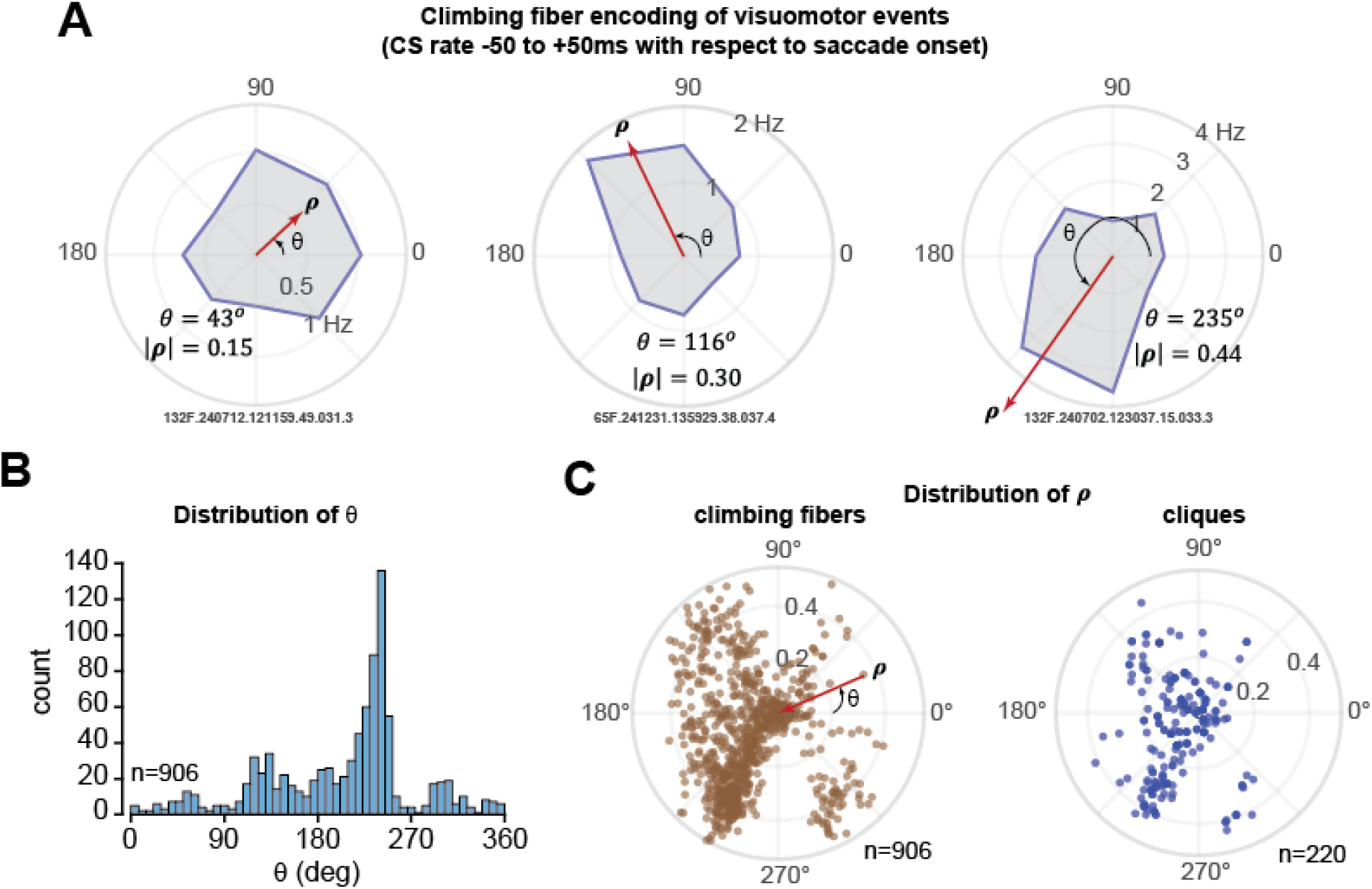
Quantifying the preferred direction of the climbing fibers. **A**. Encoding sensory events (presentation of a random target) by three climbing fibers. The CS firing rates are plotted as a function of direction of the target, and were fitted to a Von Mises distribution, producing a vector ***ρ*** with magnitude |***ρ***| in direction *θ*. **B**. Distribution of *θ*. C. Distribution of ***ρ*** across the climbing fibers, and across cliques, where a clique is a group of P-cells that are strongly connected with each other ^23^.

**Fig. S5.**
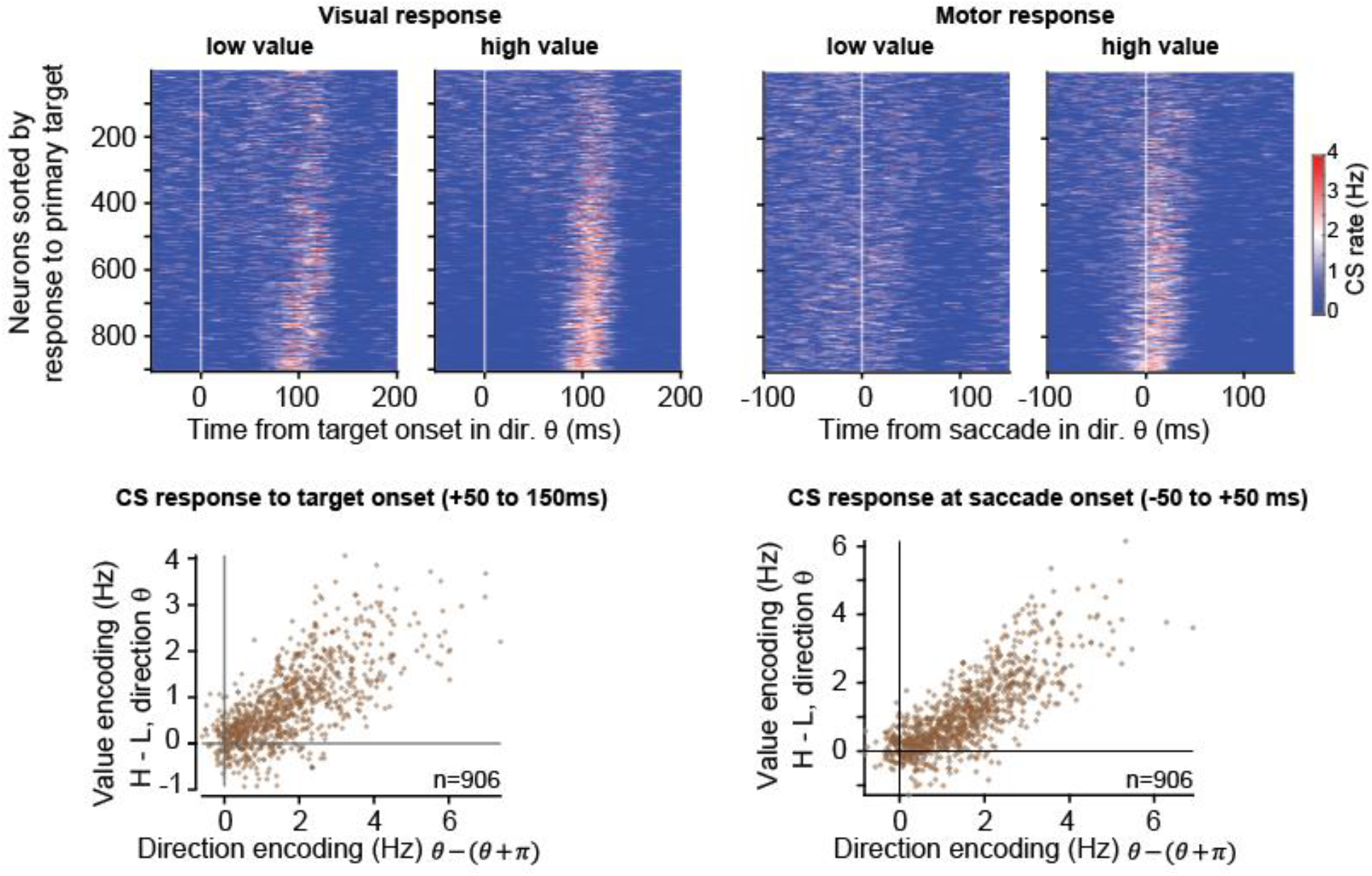
Climbing fiber encoding of direction of visuomotor event, vs. encoding of event reward value. Top row: CS rates for individual P-cells aligned to onset of the primary target, and onset of the primary saccade. Bottom row: Value encoding, measured as the difference in response for high vs. low target value, direction *θ*, plotted as a function of target or saccade direction. Individual climbing fibers encoded both the direction of the stimulus, and its value.

**Fig. S6.**
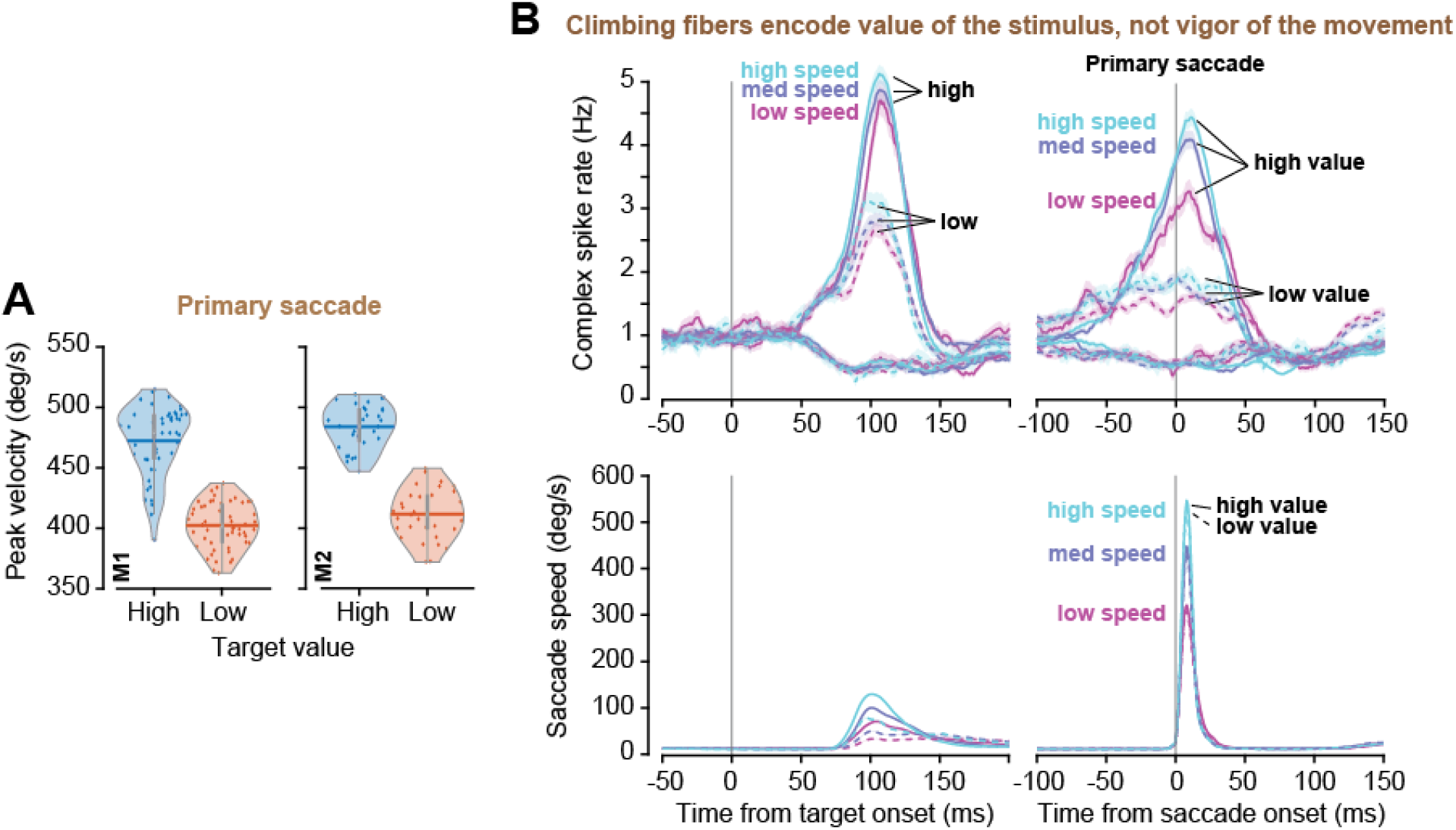
Climbing fibers principally encode stimulus reward value, not movement vigor. **A**. Peak velocity of primary saccades in the two monkeys for high and low valued targets. Each dot is the mean peak velocity for saccades in a single recording session. **B**. Complex spike firing rates aligned to target onset and saccade onset, binned by peak speed of saccades.

**Fig. S7.**
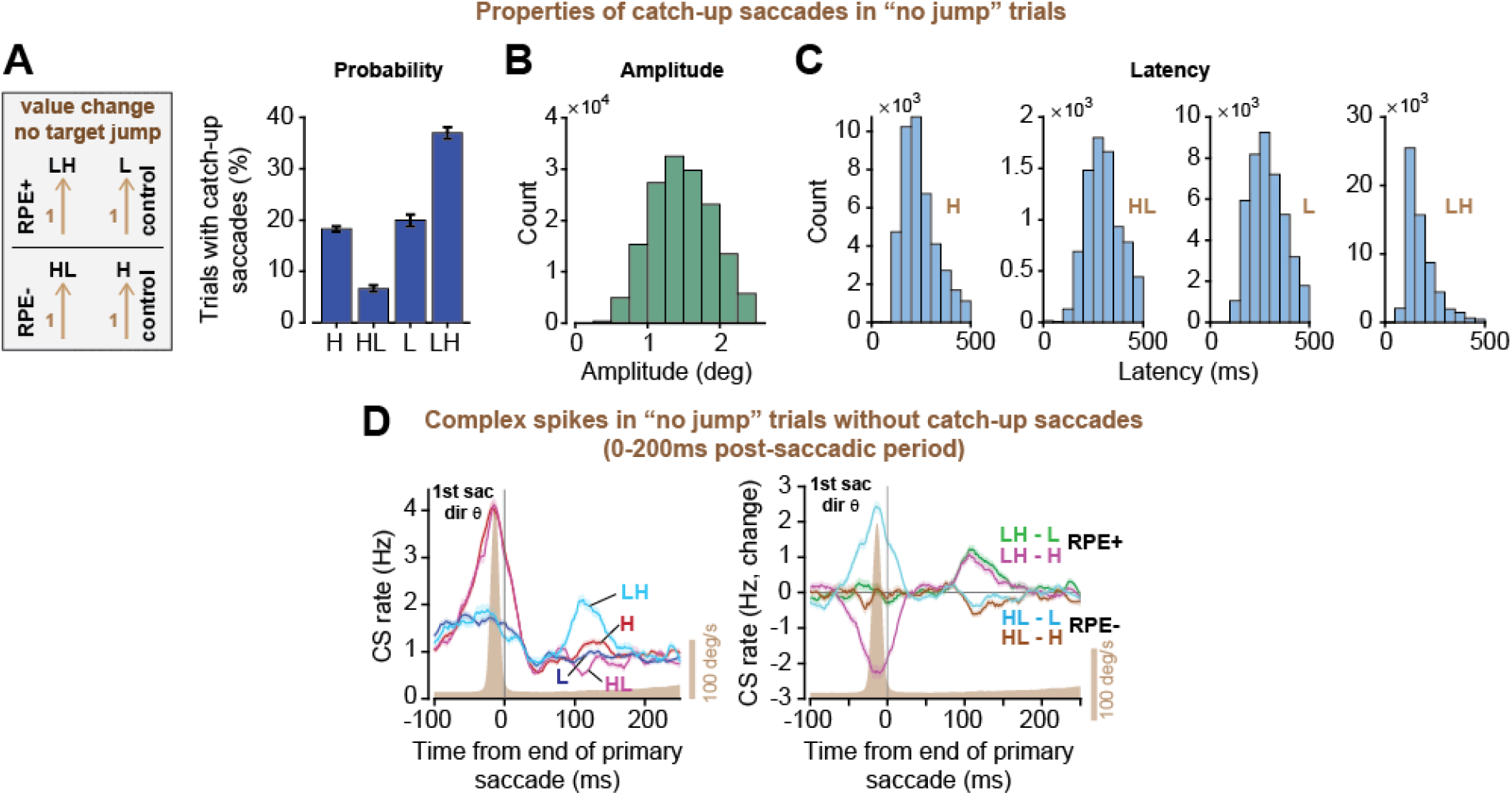
Climbing fiber activity in the trials without a target jump. **A**. Probability of catch-up saccades in no-jump trials as a function of reward context. **B**. Distribution of catch-up saccades. **C**. Latency of catch-up saccades. **D**. CS rates in no-jump trials in which catch-up saccades did not occur. Data are shown for trials in which the primary saccade was in direction *θ*.

**Fig. S8.**
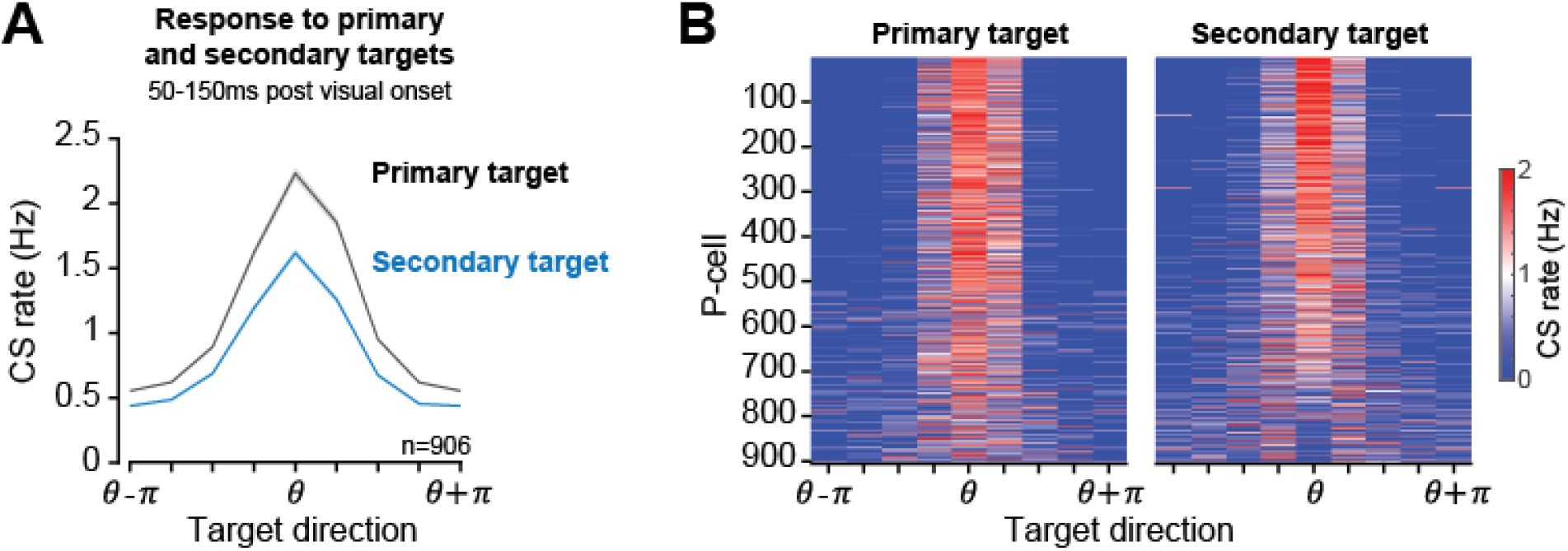
The preferred direction of climbing fibers remained consistent in response to the primary and second targets. Complex spike activity as a function of target direction was fitted to the primary target, resulting in an estimate of *θ*. The results were then used to plot the response to the secondary target. **A**. CS rates as function of target direction, 50-150ms post target onset period. **B**. Response of individual P-cells. Error bars are SEM.

## Acknowledgements

We are grateful to Mohammad Amin Fakharian, Jay Pi, Alden Shoup, Paul Hage, and Ehsan Sedaghat-Nejad, who performed the surgeries, helped with training, wrote software, and helped set up the recording chamber.

## Funding

This work was supported by grants from the National Institutes of Health (R37N128416 and R21NS145008). Hisham Elseweifi and Elijah Taeckens were supported by a training grant from the NIH (5T32EB003383).

## Author contributions

J.D. contributed to experiment design, performed software development, trained animals, collected data, analyzed data, generated figures, performed statistical testing, contributed to interpretation and project leadership. N.M. collected data, trained animals, and performed software development and data analysis. H.Y.E. designed analysis techniques, generated figures, performed statistical testing, and contributed to interpretation of results. A.D.C. designed analysis techniques, generated figures, performed statistical testing, and contributed to interpretation of results. E.A.T. contributed to interpretation of results and the theoretical framework. R.S. wrote the paper, conceived the computational framework, contributed to interpretation of results and aided project leadership.

## Notes

### Competing Interest Statement

The authors have declared no competing interest.

